# Uncovering the genetic basis of fruit volatiles in *Fragaria vesca* through GWAS reveals FvJMT2 as a methyl benzoate biosynthesis gene with insect-repellent function

**DOI:** 10.1101/2025.11.18.689018

**Authors:** Raquel Jiménez-Muñoz, Maria Urrutia, Victoriano Meco, José L. Rambla, Tuomas Toivainen, Meritxell Pérez-Hedo, José F. Sánchez-Sevilla, Alberto Urbaneja, Joaquín J. Salas, Timo Hytönen, Antonio Granell, Carmen Martín-Pizarro, David Posé

## Abstract

Strawberry aroma is a key component of fruit quality, influencing consumer preferences and playing important ecological roles, including plant defense. However, the genetic basis of volatile organic compound (VOC) biosynthesis remains only partially understood, particularly in the wild woodland strawberry *Fragaria vesca*, which has enormous potential to uncover genetic diversity within the genus for improving the commercial strawberries. Here, we performed Genome-Wide Association Studies (GWAS) across a diverse European collection of *F. vesca* accessions. We identified multiple novel candidate genes involved in the biosynthesis of diverse volatile esters, lactones, terpenoids, and methyl ketones. Among them, we characterized *FvJMT2*, a SABATH family methyltransferase which we found associated with natural variation in benzenoid esters content. Transient expression in *Nicotiana benthamiana* and strawberry fruit confirmed its role in methyl benzoate biosynthesis, while enzymatic assays demonstrated that *FvJMT2* encodes a promiscuous enzyme capable of methylating not only benzoic acid, but also cinnamic, salicylic, and jasmonic acids. Behavioral assays revealed that methyl benzoate at physiologically relevant concentrations significantly reduced the attraction of *Drosophila suzukii* flies, supporting a dual role of this VOC in both flavor and pest deterrence. Finally, natural variation analyses in wild *Fragaria* species and *F. × ananassa* cultivars showed that benzenoid esters have been largely lost in modern cultivars but retained in ancient and wild accessions. Altogether, this study provides novel insights into the genetics and ecological relevance of strawberry volatiles and identifies candidate loci and alleles for future studies.

## INTRODUCTION

Strawberry (*Fragaria* × *ananassa* Duch.) is one of the most economically important berry crops worldwide for its characteristic aroma and flavor, as well as its nutritional and health-promoting properties ^1,2^. The woodland strawberry (*Fragaria vesca* L.) is a diploid ancestor and wild relative of the octoploid cultivated strawberry. It has been traditionally employed as a model species in genetic and molecular studies and serves as a reservoir of genetic diversity for the genus. Due to its high genetic variability, *F. vesca* is a valuable resource for research aimed at improving fruit quality, yield, shelf-life, and sensory traits such as aroma and flavor, which are receiving increasing attention ^3^.

Strawberry aroma is largely determined by a subset of its volatilome profile, defined by the composition and relative abundance of key volatile organic compounds (VOCs) ^4^, which is strongly correlated with consumer preference and overall fruit liking ^3,5^. Numerous studies have characterized the volatilome of strawberry fruits in both *F.* × *ananassa* and *F. vesca*, with the latter generally exhibiting a richer and more diverse VOC profile ^4,6^. Several efforts have also addressed the genetic architecture of the cultivated strawberry volatilome, shedding light on the genetic control of biochemical pathways involved in the synthesis of key strawberry fruit volatiles, such as mesifurane, methyl anthranilate, γ-decalactone, and linalool ^7–9^. However, fewer studies have focused on woodland strawberry ^10,11^.

In addition to their sensory contributions, several VOCs commonly present in the strawberry fruit, such as terpenes and green leaf volatiles, have been associated with plant signaling and defense responses in other species ^12–15^. Methyl esters of carboxylic acids are of particular interest, as they not only contribute to fruit flavor and aroma but also function as inactive derivatives of phytohormones and as signaling molecules in plant defense ^16–18^. Among them, methyl benzoate and methyl cinnamate stand out for their dual function. Methyl benzoate is characterized by a sweet, balsamic, spicy, and heady floral odor, while methyl cinnamate carries spicy and fruity balsamic notes ^9,19,20^. Notably, both compounds have also been reported to exert insect-repellent effects, with methyl benzoate showing toxicity against various insect pests, including *Drosophila suzukii* (Matsumura) (Diptera: Drosophilidae), a major threat to soft fruit production ^20,21,22^. Females of this fruit fly use their serrated ovipositor to pierce ripening fruits and lay eggs inside the flesh ^23^, causing extensive damage and severe economic losses worldwide ^24,25^. In this context, volatile esters such as methyl benzoate and related compounds have gained increasing attention as naturally occurring bioactive molecules with both insecticidal and repellent potential ^20,26,27^.

The biosynthesis of these methyl esters is catalyzed by S-adenosyl-L-methionine (SAM)-dependent methyltransferases belonging to the SABATH family, which transfer methyl groups from SAM to low molecular weight carboxylic acids, producing small methyl esters. This family includes enzymes such as BAMT (SAM:benzoic acid carboxyl methyltransferase), CCMT (Cinnamate/p-coumarate carboxyl methyltransferases), JMT (SAM:jasmonic acid carboxyl methyltransferase), and SAMT (SAM:salicylic acid carboxyl methyltransferase) ^17,28^, many of which have been characterized in different plant species ^29–34^. In strawberry, a gene encoding a jasmonic acid methyltransferase (*FvJMT*) has been described; it is expressed in immature green fruits and capable of catalyzing the methylation of multiple carboxylic acid substrates, including jasmonic acid ^35^. In addition, another SAM dependent O-methyltransferase (*FaOMT*) has been identified as responsible for mesifurane biosynthesis in strawberry fruits ^8,36^.

Recently, a European collection of over 200 *F. vesca* accessions—part of the Eurasian subpopulation, which is the most genetically diverse within the species ^37^—was comprehensively characterized through genome resequencing and fruit volatilome profiling ^38,39^. Population structure analysis revealed two main groups corresponding to Eastern and Western Europe, along with a more complex underlying structure involving admixture among eight genetic clusters ^39^. In parallel, the volatilome of ripe fruits from this collection exhibited a broad range of profiles, with distinctive VOC signatures associated with specific geographic regions ^38^. Together, these datasets provide a unique opportunity to identify the genetic basis of natural variation in fruit aroma through Genome-Wide Association Studies (GWAS). GWAS is a powerful tool for dissecting the genetic basis of complex traits in large natural populations or germplasm collections, and can be applied here to discover genetic markers associated with VOC content and, potentially, the underlying causal genes. GWAS has been successfully applied in many horticultural crops, including tomato, Japanese pear, apple, and citrus ^40–43^. In cultivated strawberry, this approach has enabled the identification of QTLs and development of genetic markers related to resistance, agronomic performance, and fruit quality ^7,44–48^. When applied to a natural collection of a genetically diverse wild relative of a major horticultural crop, such as *F. vesca*, GWAS can facilitate the discovery of novel alleles and candidate genes for traits of agronomic interest, enabling their potential introgression into breeding programs.

Here, we provide an extensive genetic characterization of the woodland strawberry volatilome in a large European natural collection of the species. We identify stable associations between the accumulation of specific volatiles and narrow genomic regions and propose candidate genes associated with these traits. In addition, we functionally validate a novel jasmonic acid carboxylmethyltransferase in strawberry (*FvJMT2*), which is active against a variety of carboxylic acids, including jasmonic, benzoic, salicylic, and cinnamic acids. Finally, we evaluate the accumulation patterns of methyl benzoate and methyl cinnamate across different *Fragaria* species and *F.* × *ananassa* cultivars, and explored their potential ecological role in defense against *D. suzukii*.

## RESULTS

### GWAS identified known genes for VOCs biosynthesis in ripe woodland strawberry fruits

To identify genetic regions linked to the phenotypic expression of volatile profiles in ripe fruit of woodland strawberry, GWAS were performed using the available genotypic and volatilome datasets ^38,39^. The initial panel of 2.9 M SNPs was filtered to retain common variants with a minor allele frequency ≥ 5 %, resulting in a final dataset of 1.4 M SNPs. The volatilome dataset consisted of relative quantifications of 99 VOCs measured in ripe receptacles across two independent seasons, with 125 and 170 accessions analyzed, respectively. For the 113 accessions present in both seasons, data were also combined using estimated marginal means.

GWAS were then conducted independently for each harvest and for the combined dataset. A total of 312 significant associations were identified for 76 VOCs. In particular, 101 associations were found in the first harvest, 105 in the second, and 106 in the combined dataset (Table S1). Several of these associations colocalized across harvests, or between an individual harvest and the combined dataset, either for the same compound or for different compounds within the same biosynthetic pathway. These were considered stable associations and are reported in Table 1 and shown in Figure 1.

**Figure 1.**
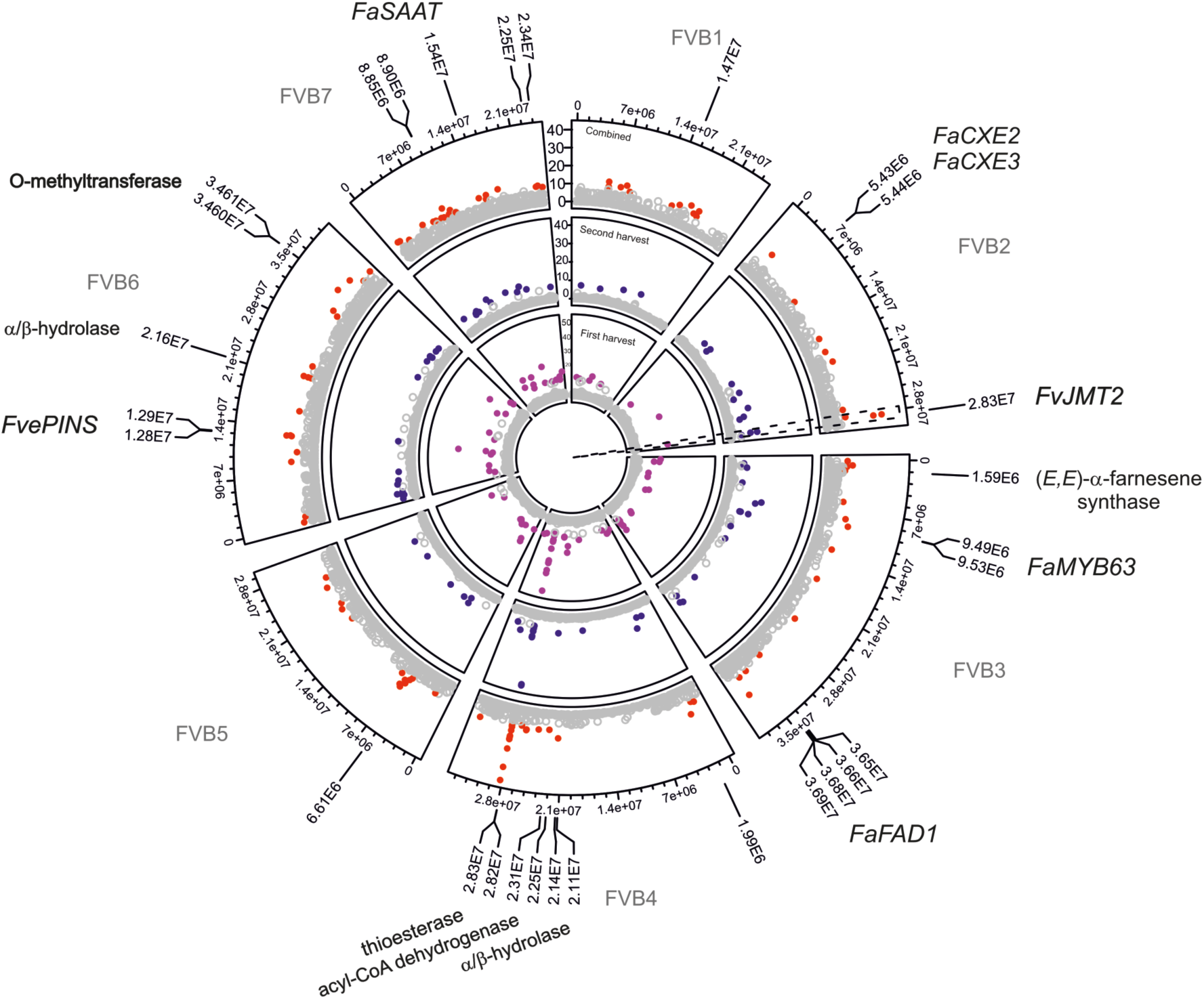
GWAS of the woodland strawberry volatilome. Circular Manhattan plots display GWAS results for the first, second, and combined harvests in the inner, middle, and outer circles, respectively. Chromosomes are arranged clockwise, with genomic coordinates indicated on the outermost circle. Pink, blue, and red dots represent significant associations between SNPs and VOCs in the first, second, or combined harvest, respectively. Genomic coordinates of stable, significant associations listed in Table 1 are marked with single slashes or brackets. Candidate gene associations previously reported are indicated in regular font, while novel associations identified in this study are shown in bold.

**Table 1.** Summary of GWAS results. Results are sorted by chromosome and show associations identified for one or more compounds within the same biosynthetic pathway, detected in at least two datasets (individual harvests and/or combined dataset). The SNP peak refers to the SNP with the lowest association p-value. The position indicates either a single SNP or a genomic interval encompassing SNP peaks identified across different harvests and/or analytical methods. Candidate genes were proposed based on colocalization with previously reported or functionally validated genes from the literature, and/or supported by gene predictions and functional annotations from the current *F. vesca* genome ^49^, together with the observed associations.

| Pathway | Metabolite | SNP peak | Stab.<sup>a</sup> | Chr. | Position | Candidate gene | Ref.<sup>b</sup> |
| --- | --- | --- | --- | --- | --- | --- | --- |
| cuticular-wax | 2-heptanol | SFVB1_14742884 | 2 | 1 | 14,742,884 | - | - |
| volatile benzenoid esters and esters | methyl-cinnamate | SFVB2_28340942 | 3 | 2 | 28,340,942 | FvH4_2g39441 (*FvJMT2*) | - |
|  | methyl-3-hydroxyoctanoate | SFVB2_28340942 | 3 |  |  |  |  |
|  | methyl benzoate | SFVB2_28340942 | 1 |  |  |  |  |
| terpenoid | (*E,E*)-α-farnesene | SFVB3_1590202 | 2 | 3 | 1,590,202 | FvH4_3g03000 ((*E,E*)-α-farnesene synthase); FvH4_3g03042 (*FaNES*) | 7 |
| volatile benzenoid | eugenol | SFVB3_9494371 | 3 | 3 | 9,494,371–9,534,713 | FvH4_3g15320 (*FaMYB63*) | 50 |
| cuticular-wax | 2-tridecanone | SFVB3_36539186 | 2 | 3 | 36,539,186–36,682,454 | - | - |
| lactones | γ-decalactone | SFVB3_36899367 | 3 | 3 | 36,899,367–36,903,084 | FvH4_3g44270 (*FaFAD1*) | 51 |
| volatile benzenoid esters | methyl cinnamate | SFVB4_1995255 | 3 | 4 | 1,995,255 | - | - |
| volatile esters | methyl butanoate | SFVB4_21171862 | 2 | 4 | 21,171,862–21,436,796 | FvH4_4g17380 ($\alpha/\beta$-hydrolase) | - |
| fatty acid degradation | 1-penten-3-ol | SFVB4_22521848 | 2 | 4 | 22,521,848–23,159,434 | FvH4_4g18700 (acyl-CoA dehydrogenase) | - |
| cuticular-wax and volatile esters | 2-undecanol | SFVB4_28255561 | 3 | 4 | 28,255,561–28,276,260 | FvH4_4g27130; FvH4_4g27160 (thioesterases) | - |
|  | 2-undecanone | SFVB4_28255561 | 3 |  |  |  |  |
|  | 2-heptanol | SFVB4_28263557 | 1 |  |  |  |  |
|  | 2-nonanone | SFVB4_28268200 | 3 |  |  |  |  |
|  | 2-nonanol | SFVB4_28268200 | 2 |  |  |  |  |
|  | 1-methyloctyl-butanoate | SFVB4_28268200 | 3 |  |  |  |  |
|  | ethyl dodecanoate | SFVB4_28269967 | 2 |  |  |  |  |
|  | methyl dodecanoate | SFVB4_28269967 | 3 |  |  |  |  |
|  | 2-heptanone | SFVB4_28273119 | 3 |  |  |  |  |
|  | methyl decanoate | SFVB4_28275857 | 2 |  |  |  |  |
| not assigned | acetophenone | SFVB5_6607454 | 2 | 5 | 6,607,454 | - | - |
| terpenoid | α-pinene | SFVB6_12829851 | 2 | 6 | 12,785,984–12,879,960 | FvH4_6g19300 (*FvePINs*) | 52 |
| | α-terpineol | SFVB6_12829851 | 2 | 6 | | | |
| volatile esters | butyl butanoate | SFVB6_21571423 | 2 | 6 | 21,571,423 | FvH4_6g28031 ($\alpha/\beta$-hydrolase) | - |
|  | hexyl butanoate | SFVB6_21571423 | 1 | 6 |  |  |  |
|  | propyl butanoate | SFVB6_21571423 | 1 | 6 |  |  |  |
| furanes | mesifurane | SFVB6_34607987 | 3 | 6 | 34,607,987–34,646,690 | FvH4_6g45070 (O-methyltransferase) | - |
| volatile esters | methyl dodecanoate | SFVB7_15462516 | 3 | 7 | 15,459,398–15,464,951 | FvH4_7g18570 (*FaSAAT*) | 53 |
|  | methyl decanoate | SFVB7_15462018 | 1 | 7 |  |  |  |
|  | ethyl dodecanoate | SFVB7_15459398 | 1 | 7 |  |  |  |
| terpenoid | linalool | SFVB7_22529616 | 2 | 7 | 22,529,616–23,439,680 | - | - |

<sup>a</sup>Stab. Number of analyses in which the association was found.
<sup>b</sup>Ref. Bibliographical reference reporting candidate gene.

Importantly, our results validated previously reported QTL associations and biosynthetic genes known to regulate various well-characterized strawberry volatiles. Among these, a stable association was identified between γ- decalactone levels and SNPs within a 3.7-kb region on chromosome 3 (Chr3: 36,899,367–36,903,084 bp), located in the promoter of FvH4_3g44270, the *F. vesca* ortholog of *FaFAD1* (Tables 1 and S1; Fig. S1A). This gene encodes a fatty acid desaturase involved in γ-decalactone synthesis during *F.* × *ananassa* fruit ripening, which contributes to the characteristic peach-like aroma ^51,54^.

For the phenylpropanoid derivative eugenol, an important odor-active compound present at lower levels in octoploid strawberry compared to *F. vesca* ^4,9,55^, we found a 40.3-kb region on chromosome 3 (Chr3: 9,494,371–9,534,713) containing FvH4_3g15320, the *F. vesca* ortholog of *FaMYB63* (Tables 1 and S1; Fig. S1B). This R2R3-type MYB transcription factor, like *FaEOBII,* has been shown to activate key genes in the eugenol biosynthetic pathway, such as *EUGENOL SYNTHASE 1* and *2* (*FaEGS1* and *FaEGS*2), and *CINNAMYL ALCOHOL DEHYDROGENASE1* (*FaCAD1*) ^50,56,57^.

Volatile esters such as methyl decanoate, methyl dodecanoate, and ethyl dodecanoate were associated with a region on chromosome 7 (Chr7: 15,459,398–15,464,921) (Tables 1 and S1; Fig. S1C) near FvH4_7g18570, the *F. vesca* ortholog of *FaSAAT*, a well-known alcohol acyltransferase involved in ester biosynthesis during fruit ripening (Aharoni et al. 2000).

A strong association was also detected for the terpenoid (*E,E*)-α- farnesene, with a SNP located on chromosome 3 (Chr3: 1,590,202 bp) in the promoter region of FvH4_3g03000 (Tables 1 and S1; Fig. S1D), which encodes a terpene synthase orthologous to the putative (*E,E*)-α-farnesene synthase previously identified by Barbey et al. ^7^. This region is also near FvH4_3g03042 and FvH4_3g03081, orthologs of *NEROLIDOL SYNTHASE* genes (*FaNESs*), which were previously associated with the terpenoid synthesis (linalool and nerolidol) in the same study ^7^.

Additionally, a significant association was found in the second harvest between (*E*)-2-hexenal and a SNP located on chromosome 2 (Chr2: 5,203,012) (Table S1; Fig. S1E), positioned 229 kb upstream of carboxylesterases FvH4_2g06540 and FvH4_2g06560 (orthologs of *FaCXE3* and *FaCXE2*, respectively) which catalyze the hydrolysis of (*E*)-2-hexenyl acetate to (*E*)-2- hexenol which in turn can be further converted to (*E*)-2-hexenal by an alcohol dehydrogenase ^58^.

Thus, the co-localization of these SNPs with previously identified biosynthetic genes for VOCs such eugenol, volatile esters, and (*E*,*E*)-α- farnesene, among others, in both *F.* × *ananassa* and *F. vesca*, highlights the robustness of our GWAS approach in capturing genetic determinants of volatile biosynthesis and supports the overall reliability of our analyses.

### GWAS identified novel candidate genes for VOCs biosynthesis in ripe woodland strawberry fruits

In addition to identifying SNPs linked to previously validated or proposed candidate genes, our analysis uncovered novel associations with VOCs of interest, either due to their significant contribution to fruit aroma and/or their potential biological function (Fig. 1). Among these, stable associations were also identified for the terpenoids α-pinene and α-terpineol, with SNPs mapping to a region on chromosome 6 containing FvH4_6g19300, a gene annotated as an α- pinene synthase (*FvePINS*) (Tables 1 and S1; Fig. S1F).

A significant, novel association peak for mesifurane was identified on chromosome 6 (Chr. 6: 34,607,987–34,646,690 bp) (Table 1 and S1; Fig. S1G), in a region containing FvH4_6g45070, an O-methyltransferase-encoding gene distinct from FvH4_7g32990, the *F. vesca* ortholog of *FaOMT*, which was previously reported to regulate the content of this VOC in *F.* × *ananassa* ^7,8,36^, and is therefore a candidate gene potentially involved in the biosynthesis of this VOC in *F. vesca*.

In addition, robust and stable association for methylketones, as well as their derived secondary alcohols and precursor esters, was detected within a 20.7 kb genomic region on chromosome 4 (Chr4: 28,255,561–28,276,260 bp) (Tables 1 and S1; Figs. S1C and H). This region primarily spans the intergenic region between two thioesterase-encoding genes, whose enzymatic activities have been implicated in the biosynthesis of these compounds in tomato ^59^.

Furthermore, an association cluster for short-chain butanoic acid esters (propyl-, butyl-, and hexyl-butanoate) was identified on chromosome 6 (Chr. 6: 21,571,423 bp) (Tables 1 and S1; Fig S1I), adjacent to gene FvH4_6g28031, annotated as a putative α/β-hydrolase. Additionally, a stable association was found for the methyl ester of butanoic acid (methyl butanoate) on chromosome 4 (Chr. 4: 21,171,862–21,436,796 bp), which includes the gene FvH4_4g17380, another α/β-hydrolase. These proteins belong to a family of enzymes that catalyze the formation and cleavage of ester bonds ^60^ and therefore emerge as strong candidates for involvement in ester biosynthesis.

Finally, the fatty acid degradation-derived volatile 1-penten-3-ol was significantly associated with a region spanning chromosome 4 (Chr. 4: 22,521,848–23,159,434 bp) (Tables 1 and S1; Fig. S1J), which includes the gene FvH4_4g18700, annotated as an acyl-CoA dehydrogenase (ACAD). This class of enzymes catalyzes the initial step of each cycle of mitochondrial fatty acid β- oxidation ^61^ and thus represents a potential candidate gene for mediating 1- penten-3-ol release.

Altogether, our GWAS analysis has enabled the identification of novel candidate genes and genomic loci involved in volatile biosynthesis, offering new targets for further functional studies and aroma-focused breeding strategies in strawberry.

### Identification of *FvJMT2* as a candidate gene for the biosynthesis of volatile benzenoid esters

Among the novel associations identified in our GWAS was a SNP located on chromosome 2 (Chr. 2: 28,340,942) (Tables 1 and S1; Figs. 1 and S1K), linked to the accumulation of the volatile benzenoid esters methyl benzoate and methyl cinnamate. This SNP (SFVB2_28340942) lies within the coding region of FvH4_2g39441, which encodes an SAM:jasmonic acid carboxylmethyltransferase. Unlike in our GWAS analysis, the *F.* × *ananassa* ortholog of this gene has been associated with variation in methyl anthranilate (methyl 2-aminobenzoate) content and described as *FanAAMT-like* ^7^ (hereafter referred to as *FvJMT2*). Notably, the SNP is located 275 nucleotides downstream of the *FvJMT2* start codon and results in a cytosine-to-adenosine substitution (C275A) (Fig. 2A). While the *FvJMT2^C^*^275^ allele encodes the full-length predicted protein of 390 amino acids, the *FvJMT2^A^*^275^ allele introduces a premature stop codon (Ser92Stop), resulting in a truncated protein of only 91 amino acids (Fig. S2). Interestingly, the full-length *FvJMT2^C^*^275^ allele is prevalent in the population, while the truncated *FvJMT2^A^*^275^ allele exhibits a minor allele frequency of 12% and is present in samples collected from Iceland, northern Norway, United Kingdom and Portugal (Table S2).

**Figure 2.**
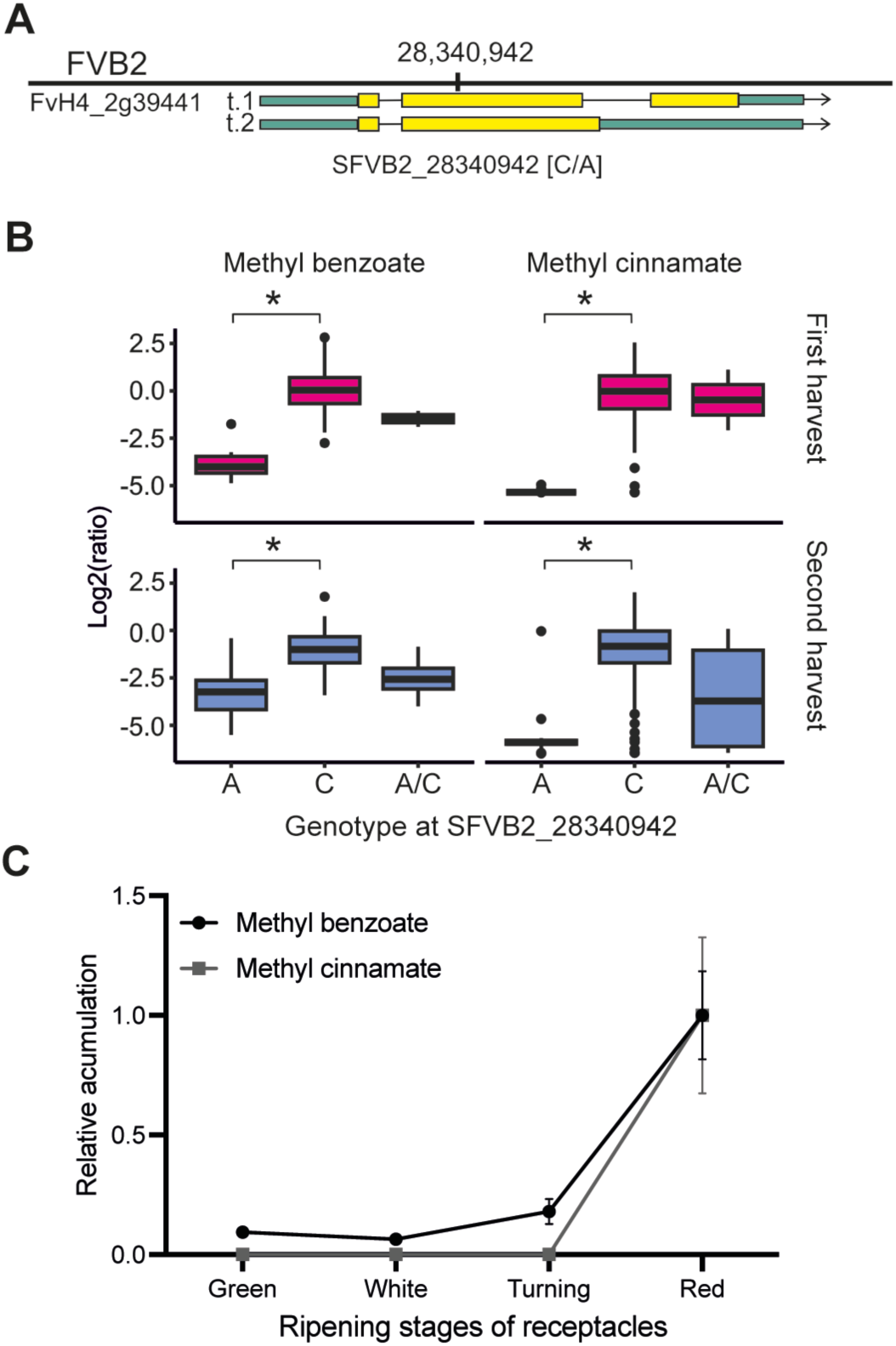
*FvJMT2* is a candidate gene for the biosynthesis of methyl benzoate and methyl cinnamate. **A)** Genomic region surrounding the candidate SNP SFVB2_28340942, located in the second exon of FvH4_2g39441. **B)** Relative accumulation of methyl benzoate and methyl cinnamate according to the genotype at SNP SFVB2_28340942 genotype. Genotypes are indicated using single-letter codes: homozygous adenosine (A), homozygous cytosine (C), and heterozygous (A/C). Asterisk denote significant differences in compound accumulation (Student’s *t*-test, *p* < 0.01). **C)** Relative content of methyl benzoate and methyl cinnamate in fruit receptacles of *F. vesca* ‘Reine des Vallées’ at four developmental stages. Quantification is relative to the red stage.

Woodland strawberry fruits from accessions carrying the truncated *FvJMT2^A2^*^75^ allele accumulated significantly lower levels of methyl benzoate and methyl cinnamate, whereas heterozygous accessions displayed intermediate levels of these volatiles (Fig. 2B). These findings support the hypothesis that a functional full-length FvJMT2 protein contributes to the biosynthesis of these compounds.

To further investigate the putative function of FvJMT2, we performed a sequence homology analysis using BLASTp, which revealed high similarity to jasmonic acid carboxyl methyltransferases (JMTs) from the *Rosaceae* family. These enzymes belong to the SABATH family of SAM-dependent methyltransferases, which catalyze the methylation of a broad range of small carboxylic acids, including benzoic and cinnamic acids, as well as jasmonic and salicylic acids, among others ^62^.

We then compared the FvJMT2 protein sequence with those of functionally characterized enzymes, including JMT, BAMT, SAMT, BSMT, AAMT, and CCMT from various plant species ^17^, including *CbSAMT* from *Clarkia breweri*, the first structurally resolved SABATH enzyme ^63^ (Fig. S3). Using CbSAMT as a reference, alignment analysis showed that FvJMT2 conserves all four SAM binding sites but exhibits semi- and non-conservative substitutions at two substrate-binding positions, specifically Y166S and I250H (Fig. S3). These substitutions are shared by all JMTs included in the analysis, with which FvJMT2 shares high sequence similarity (60–76%). Interestingly, CmBAMT from melon exhibited greater similarity at the substrate binding sites with JMTs than with other BAMTs. Consistent with these findings, a neighbor-joining consensus tree grouped FvJMT2 with other JMTs and CmBAMT (Fig. S4), supporting a closer evolutionary relationship among these enzymes and reinforcing its functional classification as a JMT-like member of the SABATH methyltransferase family.

### *FvJMT2* expression and the content of its associated VOCs increase during strawberry fruit ripening

Next, we analyzed the content of *FvJMT2*-associated VOCs, i.e., methyl benzoate and methyl cinnamate, by GC-MS in receptacles of *F. vesca ssp. vesca* cv. ‘Reine des Vallées’ (C275 allele) at four ripening stages (green, white, turning, and red). This analysis revealed that the accumulation of these VOCs increased in receptacles during fruit ripening, reaching their highest levels at the ripe stage (Fig. 2C). In particular, methyl benzoate content was more than 5-fold higher in ripe receptacles compared to earlier developmental stages (green, white, and turning), while methyl cinnamate and methyl 3-hydroxyoctanoate were undetectable at these earlier stages (Fig. 2C). Interestingly, this accumulation pattern correlated with the transcriptional upregulation of *FvJMT2* during fruit ripening, as previously reported in the accession ‘Yellow Wonder 5AF7‘ (also carrying the C275 allele) ^64,65^ (Fig. S5). A similar expression pattern was observed for its ortholog in *F.* × *ananassa* cv. Camarosa (*FaJMT2*; FxaC_8g06390), whose expression also increased during receptacle ripening ^66,67^ (Fig. S5). Thus, the correlation between *FvJMT2* expression and the accumulation of its associated VOCs further supports its role in their biosynthesis during strawberry fruit ripening.

### Heterologous and homologous transient overexpression of *FvJMT2* leads to methyl benzoate accumulation

To functionally validate the role of FvJMT2 in the biosynthesis of volatile benzenoid esters, we performed both heterologous and homologous transient overexpression experiments in *Nicotiana benthamiana* leaves and *F. vesca* receptacles, respectively. Specifically, constructs carrying the two allelic variants of *FvJMT2* were generated: 1) *35S:FvJMT2^C2^*^75^, containing the full CDS (1,173 nt) encoding the complete FvJMT2 protein; and 2) *35S:FvJMT2^A2^*^75^, also containing the full CDS but encoding the truncated protein due to the premature stop codon. Heterologous overexpression in *N. benthamiana* leaves was first confirmed by qPCR. Notably, transcript levels of *FvJMT2^A275^* were significantly lower than those of *FvJMT2^C275^* (Fig. 3A), likely due to nonsense-mediated mRNA decay triggered by the premature stop codon ^68^. To avoid this issue, we generated an additional construct (*35S:FvJMT2^A2^*^75^^Δ^*^C^*) containing only the first 275 nt of the CDS, spanning from the start codon to the premature stop codon. Consistent with transcript instability of the full *FvJMT2^A2^*^75^ transcript, this truncated version showed expression levels comparable to *FvJMT2^C275^* (Fig. 3A). These samples were then analyzed for benzenoid esters content by GC-MS. Only overexpression of the full-length *FvJMT2^C275^* allele led to a significant increase in methyl benzoate levels. In contrast, overexpression of *FvJMT2^A275^*^Δ^*^C^* did not result in methyl benzoate accumulation, with levels comparable to the mock control (Fig. 3B). Interestingly, methyl cinnamate was not detected in this heterologous assay, possibly due to the absence of its precursor, cinnamic acid, in *N. benthamiana* leaves and/or a higher catalytic preference of FvJMT2 for benzoic acid over cinnamic acid. Together, these results support that the C275 allele of FvJMT2 encodes a functional protein involved in methyl benzoate biosynthesis.

**Figure 3.**
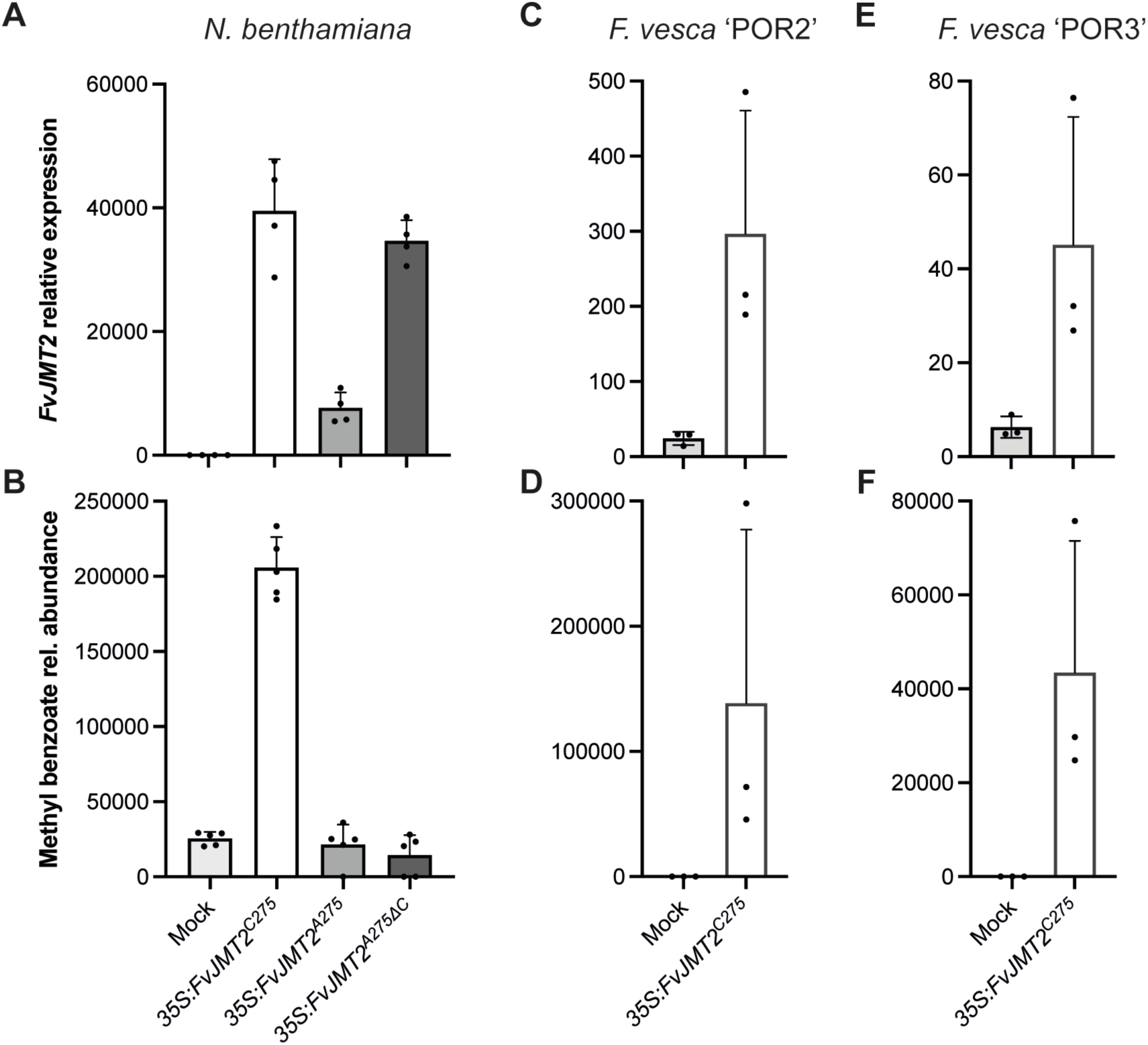
Transient overexpression of *FvJMT2* in *Nicotiana benthamiana* leaves and *Fragaria vesca* fruits. **A)** Relative expression of *FvJMT2* constructs in *N. benthamiana* leaves, measured by quantitative PCR. **B)** Relative methyl benzoate content in *N. benthamiana* leaves agroinfiltrated with *FvJMT2* constructs. **C** to **F)** Relative expression of *FvJMT2* (**C**, **E**) and methyl benzoate content (**D**, **F**) in fruits of *F. vesca* accessions ‘POR2’ (**C**, **D**) and ‘POR3’ (**E**, **F**) agroinfiltrated with *FvJMT2^C275^*.

To further confirm this activity in a homologous system, the functional *FvJMT2^C275^*allele was transiently overexpressed in *F. vesca* fruit receptacles (Fig. 3C, E) from two accessions, ‘POR2’ and ‘POR3’, which naturally carry the non-functional A275 allele and produce low levels of methyl benzoate and methyl cinnamate, as previously described ^38,39^. Transient overexpression of *FvJMT2^C275^* in the receptacles of these accessions led to a significant increase in methyl benzoate content (Fig. 3D, F), while methyl cinnamate levels remained unchanged as in *N. benthamiana*. These findings strongly support the role of FvJMT2 in methyl benzoate biosynthesis and suggest that the enzyme may exhibit a substrate preference for benzoic acid over cinnamic acid.

### FvJMT2 enzyme activity assay reveals benzoic acid as its preferred substrate

To further characterize the biochemical properties of FvJMT2 (specifically, the functional FvJMT2^C275^ allele), we conducted *in vitro* enzymatic activity assays using a range of potential substrates. Substrate specificity was evaluated through independent assays using a diverse panel of phenolic and structurally related carboxylic acids. In addition to benzoic and cinnamic acids, the panel included jasmonic, salicylic, *p-*coumaric, feruloic, caffeic, (-)-quinic, and *p*-hydroxy phenylacetic (*p-*HPA) acids, as well as gallic acid monohydrate. As shown in Table 2, FvJMT2 exhibited the highest activity toward benzoic, salicylic, and jasmonic acids, moderate activity toward cinnamic acid, and only residual activity with the remaining substrates.

**Table 2.** FvJMT2 enzymatic activity and kinetic parameters with different substrates. Enzyme activity is reported as the average ± standard deviation (SD) (n=3), expressed in pmol of substrate consumed per minute per milligram of protein. Enzyme kinetic parameters include maximum velocity (Vmax), Michaelis constant (Km), and the catalytic efficiency (Vmax/Km).

| Substrate | Activity |  | Kinetic parameters |  |  |
| --- | --- | --- | --- | --- | --- |
|  | average | SD | Vmax | Km | Vmax/Km |
| | pmol min<sup>-1</sup> mg<sup>-1</sup> | | pmol min<sup>-1</sup> mg<sup>-1</sup> | mM | μL min<sup>-1</sup> mg<sup>-1</sup> |
| Benzoic acid | 4,365.70 | 37.99 | 5,476.25 | 0.577 | 9.49 |
| Cinnamic acid | 85.67 | 1.56 | 145.10 | 0.597 | 0.24 |
| Jasmonic acid | 426.46 | 41.61 | 1,466.92 | 0.174 | 8.43 |
| Salicylic acid | 306.89 | 9.20 | 794.08 | 0.690 | 1.15 |
| *p*-coumaric acid | 8.68 | 0.38 | - | - | - |
| Feruloic acid | 6.50 | 1.16 | - | - | - |
| *p*-hpa | 5.11 | 1.92 | - | - | - |
| Caffeic acid | 4.18 | 1.30 | - | - | - |
| (-)-Quinic acid | 3.44 | 0.76 | - | - | - |
| Gallic acid monohydrate | 3.17 | 0.64 | - | - | - |

These results indicate that FvJMT2 is a substrate-promiscuous SAM-dependent methyltransferase, capable of catalyzing the methylation of multiple small carboxylic acids with variable efficiencies. To quantify its substrate preference, we performed kinetic analyses on the acids for which FvJMT2 exhibited the highest activity (Table 2, Fig. S6). Notably, and consistent with our transient overexpression results in *N. benthamiana* leaves and *F. vesca* receptacles, FvJMT2 showed the highest apparent catalytic efficiency toward benzoic acid (Vmax/Km = 9.49 μL min^-1^ mg^-1^). This was followed by jasmonic acid (Vmax/Km = 8.43 μL min^-1^ mg^-1^), for which the enzyme exhibited the highest substrate affinity (Km = 0.174 mM). Although catalytic efficiency toward salicylic acid was also relatively high, it was approximately 8-fold lower than that for benzoic acid, while activity toward cinnamic acid was markedly lower (∼40-fold lower than for benzoic acid). These results provide a biochemical explanation for the selective accumulation of methyl benzoate over methyl cinnamate observed in our *in vivo* experiments. Moreover, the significant catalytic activity toward jasmonic and salicylic acids suggests that FvJMT2 may also participate in the biosynthesis of methyl jasmonate and methyl salicylate, two well-known defense-related methyl esters ^69,70^.

### Methyl benzoate acts as a repellent VOC against the pest *Drosophila suzukii*

Given the global importance of *D. suzukii* as a pest and the previously reported insecticidal activity of volatile esters such as methyl benzoate, we next aimed to investigate whether this compound also exerts a repellent effect on *D. suzukii* in the context of strawberry fruit and to assess the potential effect of methyl cinnamate. To this end, the behavioral response of *D. suzukii* females to volatile- supplemented strawberry purée was evaluated using a two-choice arena assay, in which methyl benzoate or methyl cinnamate were applied at physiological concentrations found in ripe *F. vesca* fruits ^38^. As shown in Figure 4A, the proportion of females found in vials containing methyl benzoate was significantly lower than the expected value under a neutral preference scenario (mean = 34% ± 5; *χ*^2^ = 8.576, *p* = 0.0034), indicating a repellent effect of this compound. In contrast, no significant deviation from neutrality was observed for methyl cinnamate (mean = 46% ± 5; *χ*^2^= 0.495, *p* = 0.4817), suggesting that this compound did not elicit a directional behavioral response under the tested conditions. These findings suggest that methyl benzoate, but not methyl cinnamate, contributes not only to strawberry fruit aroma but also provides ecological advantages by exhibiting both insecticidal and potential repellent effects against *D. suzukii*.

**Figure 4.**
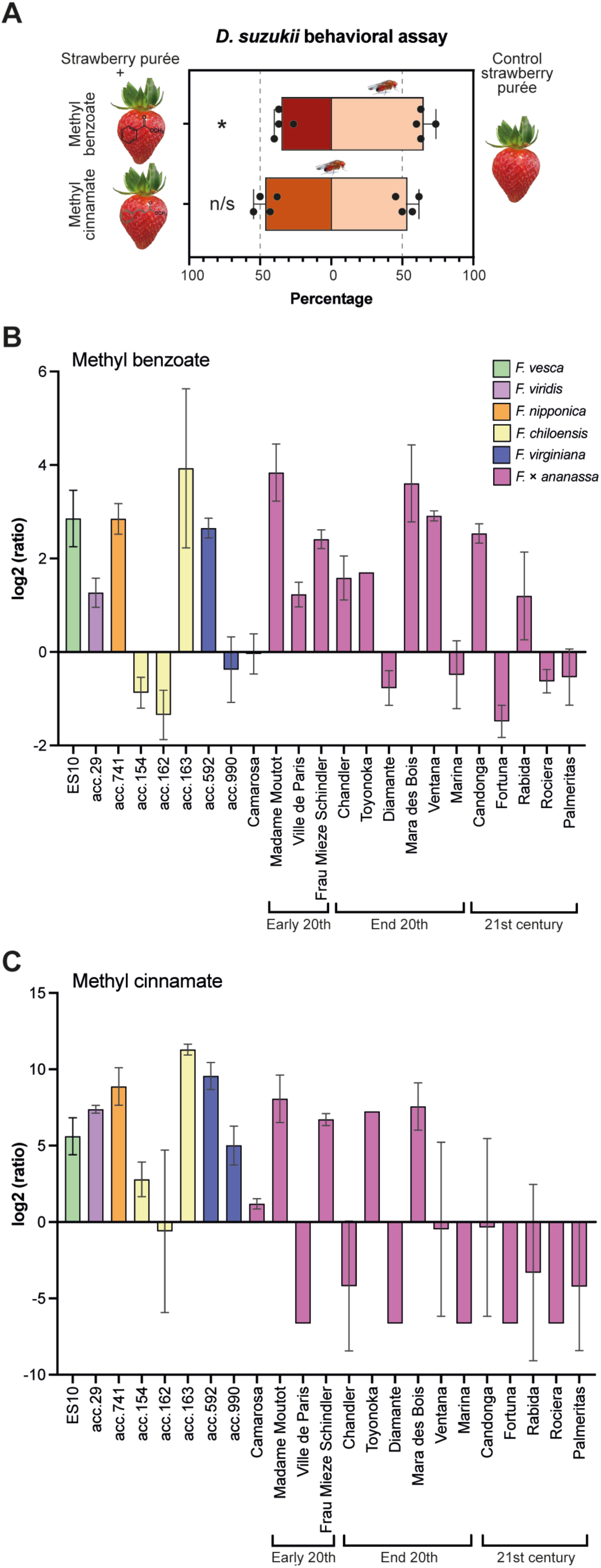
Behavioral assays of *Drosophila suzukii* using methyl benzoate and cinnamate and natural variation in their accumulation across strawberry species and cultivars. **A)** Behavioral response of *D. suzukii* females to methyl benzoate in two-choice arena assays. The graph shows the percentage of responding females found either in vials containing strawberry purée supplemented with the volatile compound (methyl benzoate or methyl cinnamate) or in control vials (n = 4 biological replicates per compound; number of individuals per replicate: 19–28). Each dot represents an independent biological replicate. Error bars indicate the standard error of the mean (SEM). The dotted grey line at 50% indicates no preference (i.e., a neutral choice). Asterisk denote statistically significant differences from 50% (*χ*^2^ tests; *p* < 0.05); n/s: not significant. **B, C)** Relative quantification of methyl benzoate (B) and methyl cinnamate (C) accumulation in ripe fruit receptacles from various *Fragaria* accessions, including diploid wild (*F. vesca*, *F. viridis*, *F. nipponica*), octoploid wild (*F. chiloensis*, *F. virginiana*), and cultivated octoploid (*F. × ananassa*) species. Values are expressed as log2 fold-change relative to the average content in *F.* × *ananassa* cv. ‘Camarosa’.

### Natural variation for volatile benzenoid esters content in Fragaria spp. And F. × ananassa cultivars

Finally, we aimed to explore the natural variation in the accumulation of both methyl benzoate and methyl cinnamate in ripe fruit receptacles across several *Fragaria* species, including the diploid species *F. vesca*, *F. viridis*, and *F. nipponica*, and the octoploids *F. chiloensis* and *F. virginiana*, which hybridized to form the cultivated species *F.* × *ananassa.* In addition, fifteen ancient and modern

### *F.* × *ananassa* cultivars were included in the analysis

Our results revealed that the content of these two VOCs varied widely among accessions, showing a generally similar pattern of accumulation with some differences (Fig. 4B, C). Methyl benzoate was highly accumulated in most wild diploid and octoploid accessions (with the exception of two *F. chiloensis* accessions and one from *F. virginiana*) and in ancient *F.* × *ananassa* cultivars, especially in ‘Madame Moutot’ and ‘Frau Mieze Schindler’, as well as in more modern cultivars, such as ‘Mara des bois’, ‘Ventana’, and ‘Candonga’, a variety bred in the 21st century. Methyl cinnamate followed a similar accumulation pattern, with generally high levels in wild *Fragaria* accessions and in some 20^th^- century cultivars; however, it showed a greater depletion in the most modern cultivars.

These results are consistent with previous studies indicating that methyl cinnamate, a key contributor to strawberry aroma, is typically abundant in wild species but often absent in modern cultivars ^3,6^. In contrast, methyl benzoate appears to be retained in some modern commercial varieties. This difference in the content of these two VOCs might contribute to the lower aromatic attributes of some modern varieties.

## DISCUSSION

In this study, we leveraged a well-characterized European collection of *F. vesca* accessions with extensive volatile and genomic data to explore natural variation in fruit VOCs and uncover their underlying genetic determinants. Our GWAS successfully recovered loci previously associated with key strawberry volatiles in *F. × ananassa*, including eugenol, methyl decanoate, (E,E)-α-farnesene, and (E)- 2-hexenal ^7,11,50,51,53^, confirming the robustness of our approach and underscoring the transferability of genetic insights across strawberry species. More importantly, we identified novel associations and proposed promising candidate genes for VOCs not previously linked to genetic variation in *F. vesca*, broadening our understanding of the molecular basis of fruit aroma.

### Novel candidate genes for volatile content in *F. vesca*

Our analysis pinpointed several new candidate genes involved in the accumulation of esters, lactones, and terpenoids, important contributors to strawberry aroma ^3,16^ . For mesifurane, we identified an O-methyltransferase as a potential regulator of its biosynthesis, distinct from the previously described *FanOMT1* ^8^.

In relation to terpenoids, the α-pinene synthase gene *FvePINS* (FvH4_6g19300) emerged as a strong candidate for the biosynthesis of both α- pinene and α-terpineol. This gene was previously reported to be upregulated during ripening in the *F. vesca* accessions ‘Hawaii4’ and ‘Ruegen’ ^52^ and shares a high 99% sequence identity with *FvPINS* (FvH4_1g05400), another α-pinene synthase-encoding gene, known to catalyze the biosynthesis of α-pinene, β- phellandrene, and β-myrcene ^71^. Although *FvPINS* is expressed in some *F. vesca* accessions ^71^, its expression is nearly absent in ‘Hawaii4’ and ‘Ruegen’ ^52^. Therefore, *FvePINS* is likely contributing to α-pinene and α-terpineol production in these and potentially other accessions. Additionally, two α/β-hydrolases, which can act as esterases ^60^, are proposed as candidate genes for butanoic acid esters accumulation.

Beyond aroma-related VOCs, we identified candidate genes for compounds potentially involved in ecological interactions. Methyl ketones, derived from fatty-acid intermediates and known for their insect-repellent activity in tomato (*Solanum lycopersicum*) ^72^, were linked to two thioesterases, which we propose as candidate genes as they encode for enzymes involved in the cleavage of thioester bonds in fatty acyl-CoA intermediates within the fatty acid pathway ^73^, which serve as precursors for methyl ketone biosynthesis. In addition, 1-penten-3-ol, a known plant defense signal ^74^, was associated with an acyl-CoA dehydrogenase, an enzyme catalyzing the initial step of fatty acid β-oxidation ^61^, and therefore a candidate for the biosynthesis of this VOC. These findings provide new insights into the genetic regulation of volatiles with ecological functions in *F. vesca*, including potential roles in biotic stress and plant-insect interactions.

### FvJMT2 participates in methyl benzoate biosynthesis

Our GWAS also identified a SNP in the coding sequence of the methyl transferase FvJMT2 (FvH4_2g39441) associated with variability in the content of the benzenoid esters methyl benzoate and methyl cinnamate, as well as the volatile fatty acid ester methyl 3-hydroxyoctanoate. The involvement of FvJMT2 was functionally validated for methyl benzoate (Fig. 3).

Remarkably, FvJMT2, previously referred to as FvCMT (carboxyl methyltransferase) by Preuß et al. ^35^, was dismissed as a non-functional enzyme. This likely resulted from a cloning error caused by a misannotation in the *F. vesca* reference genome used in that study (v1.1) ^75^. Specifically, a 24 nt region at the 3’ end led to *FvCMT* being incorrectly annotated as a gene fusion with its downstream neighbor gene, *FvJMT* (FvH4_2g39450), also reported as a functional jasmonic acid carboxyl methyltransferase ^35^. In contrast, in the present study, we functionally characterized the proper FvH4_2g39441 coding sequence, demonstrating its methyltransferase activity on diverse substrates.

Interestingly, the putative orthologs of *FvJMT2* in *F.* × *ananassa*, the *ANTHRANILIC ACID METHYL TRANSFERASE-like* (*FanAAMT-like*) genes, were previously identified as candidate genes for variation in methyl anthranilate content, a grape-like aroma compound, in a segregant population of the cultivated species ^7^. These orthologs are encoded by two homologs located on the Fvb2-1 (maker-Fvb2-1-snap-gene-255.58; FxaC_7g41870) and Fvb2-3 (maker-Fvb2-3- snap-gene-33.59; FxaC_8g06390) chromosomes ^66^. In contrast, no association was found between methyl anthranilate and *FvJMT2* in this study. Whether FvJMT2 may use anthranilic acid as a substrate remains to be elucidated.

### FvJMT2 is a promiscuous methyltransferase with potential ecological role

FvJMT2 belongs to the SABATH gene family, a group of enzymes that catalyze the methylation of carboxylic acids in plants. They are classified based on their preferred substrates (e.g., JMT, BAMT, SAMT), although activity (albeit at low rates) against multiple carboxylic acids is commonly observed in these enzymes^17^. In the present study, the *in vitro* enzymatic promiscuity of FvJMT2 was demonstrated by its ability to methylate multiple substrates, including, by decreasing activity, benzoic, jasmonic, salicylic, and cinnamic acids. This functional flexibility, together with the low enzymatic activity observed with cinnamic acid may explain why, despite our GWAS found association with methyl cinnamate, its production was not detected in transient overexpression assays in both homologous and heterologous systems. Whether the downstream neighboring gene *FvJMT*, which has been reported not to use benzoic acid as a substrate ^35^, and is therefore unlikely to participate in methyl benzoate biosynthesis, might instead be responsible for the variation in methyl cinnamate levels in the *F. vesca* population remains to be elucidated.

Such promiscuity is commonly observed in plant methyltransferases, particularly in the SABATH family, which are thought to have evolved to adapt to different molecules to meet diverse physiological needs ^63^. This flexibility enables plants to produce a broad array of specialized metabolites using a limited set of enzymes, and is considered a key mechanism driving the evolution of new functions ^76^. Our findings support this view, suggesting that FvJMT2 may contribute not only to the biosynthesis of methyl benzoate (as shown here), but also to the production of other methyl esters VOCs involved in defense, such as methyl jasmonate (MeJA) and methyl salicylate ^69,70^ In fact, *JMT* (At1g19640), the closest Arabidopsis homolog of *FvJMT2*, was reported to play a key role in MeJA biosynthesis, with overexpression leading to constitutive expression of jasmonate-responsive genes and enhanced resistance to *Botrytis cinerea* ^32^. Moreover, MeJA has been proposed to be produced in response to drought stress, subsequently inducing the production of abscisic acid ^77,78^. In addition, MeJA treatment has been shown to alleviate water stress, in part by modulating metabolism to enhance stress tolerance ^79^. Thus, the potential role of FvJMT2 in contributing to MeJA biosynthesis and the response to abiotic stress deserves further investigation.

### Methyl benzoate acts as a repellent VOC against *D. suzukii*

The spotted wing drosophila, *D. suzukii*, represents one of the most destructive pests of soft fruit crops, particularly strawberries ^22,80^, causing severe economic losses worldwide. In the United States alone, annual losses are estimated to exceed 500 million USD ^24,25^, with France, Italy, Switzerland, and Spain also being heavily affected ^81^. Our behavioral assays revealed that *D. suzukii* females exhibited a significant reduction in fly preference when exposed to physiologically relevant concentrations of methyl benzoate. The potential of methyl benzoate as a bioactive agent against *D. suzukii* was previously highlighted by Feng and Zhang ^20^, who demonstrated its acute toxicity and oviposition-deterrent effects in pre-infested blueberries. At concentrations ≥1% (i.e., >10,000 µg/g FW), methyl benzoate led to complete adult mortality and absence of larval development. Similarly, Gale and collaborators ^82^ reported that field applications of methyl benzoate at doses of 2.5% to 5% (corresponding to tens of thousands of ng/g) were highly effective in reducing *D. suzukii* infestations across various fruit crops. In contrast, the concentrations used in our study (∼200 ng/g fresh weight), which reflect the natural content reported for *F. vesca* fruits, are orders of magnitude lower and unlikely to trigger toxic effects. Although the observed repellence effect (preference index ∼34%) is moderate, the fact that it was achieved at physiologically realistic levels is noteworthy. However, the effect to native emissions on intact fruits remains to be confirmed. Nevertheless, this finding suggests that selecting or breeding genotypes with naturally higher methyl benzoate production could offer dual benefits: enhancing flavor and contributing to protection against *D. suzukii*, potentially acting as a natural defense mechanism in strawberry.

While solid evidence of methyl benzoate repellency in *D. suzukii* remains limited, clear repellent and oviposition-deterrent effects have been demonstrated in other pest species, including *Bemisia tabaci* (Gennadius) (Hemiptera: Aleyrodidae), *Tetranychus urticae* Koch (Acari: Tetranychidae), *Cimex lectularius* L. (Hemiptera: Cimicidae), and *Halyomorpha halys* (Stal) (Hemiptera: Pentatomidae) ^83,84^. These findings reinforce the potential role of methyl benzoate in behavioral disruption of *D. suzukii* under natural conditions.

### Implications for breeding and molecular marker development

Our analyses of methyl benzoate and methyl cinnamate levels across a diverse panel of *Fragaria* species (including diploid and octoploid wild accessions), as well as ancient and modern *F.* × *ananassa* cultivars, revealed substantial natural variation in the accumulation of both volatiles. Notably, methyl cinnamate has previously been reported in several woodland strawberry (*F. vesca* ‘Reine des Vallées’, and ‘Yellow wonder’) and in other highly aromatic wild strawberries, such as *F. moschata* ‘Cotta’ ^6,10^, but is largely absent in most *F.* × *ananassa* modern cultivars ^6,8,9^. In this study, we confirm these findings as most diploid and octoploid wild strawberry species, as well as ancient cultivated varieties known for their outstanding organoleptic qualities, such as ‘Frau Mieze Schindler’, ‘Madame Moutot’, and ‘Mara des Bois’, accumulate high levels of methyl cinnamate, whereas modern cultivars generally show low levels. These findings underscore the potential of using these germplasm resources to reintroduce desirable volatile traits into modern cultivars.

Importantly, the identification and functional validation of *FvJMT2* as a key gene for methyl benzoate biosynthesis opens the door to the development of molecular markers for favorable alleles. Such markers could be employed in marker-assisted breeding programs aimed at enhancing both fruit aroma and potentially natural resistance to pests such as *D. suzukii.* By leveraging natural allelic variation in *FvJMT2*, it may be possible to select for genotypes with optimized production of bioactive volatiles, offering a dual benefit for fruit quality and crop protection.

## MATERIAL AND METHODS

### Plant material

The *Fragaria vesca* collection used in this study was previously sequenced and its volatilome characterized in ripe receptacles, as described by Toivainen et al.^39^ and Urrutia et al. ^38^, respectively. For the analysis of methyl benzoate and methyl cinnamate content, a diversity panel of *Fragaria spp.* accessions were selected from the strawberry germplasm bank (ESP138) at the Instituto de Investigación y Formación Agraria y Pesquera (IFAPA, Churriana, Málaga, Spain). Ripe fruits from these accessions were harvested and immediately frozen in liquid nitrogen in three independent pools, corresponding to three biological replicates. Samples were deachened, ground to a fine powder in liquid nitrogen, and stored at -80 °C until VOCs analysis. Additionally, plants from *F. vesca* accessions ‘POR2’ and ‘POR3’ were grown and maintained in a shade house at IFAPA, while ‘Reine des vallées’ plants were cultivated in a greenhouse at the Institute for Mediterranean and subtropical Horticulture “La Mayora” (IHSM – UMA-CSIC). Fruits from this latter accession were harvested at four ripening stages: green, white, turning, and red for VOC analyses. Sample processing was performed as described for the diversity panel.

### Genome-Wide Association Study (GWAS)

GWAS analysis was performed using the volatilome and genotypic data previously generated and deposited in public repositories ^39,85^. Analyses were conducted independently on volatilome data from two harvest seasons (using average values per accession per harvest) and on the combined volatilome dataset (using estimated marginal means per genotype). Genotypic data were filtered to retain only non-redundant SNPs with a minor allele frequency (MAF) ≥ 5%. GWAS was conducted using the R package GAPIT (version 3) ^86^, applying both Multiple Loci Mixed Model (MLMM) and the Bayesian-information and Linkage-disequilibrium Iteratively Nested Keyway (BLINK) models. The first three principal components were included as covariates to account for population structure ^87,88^. Significant associations were identified using a Bonferroni corrected p-value threshold < 6.75E-9. Significant SNPs located within <1Mb of each other were considered part of the same QTL. MLMM and BLINK models were applied separately to the datasets from each harvest season, whereas only the BLINK model was used for the combined dataset.

### Bioinformatic sequence analysis

Multiple sequence alignment, sequence distance matrix, and consensus tree were performed with Geneious v.2025.1 (Biomatters, Auckland, New Zealand) using Geneious global alignment with free end gaps and Blosum62 penalty matrix for the alignment and Jukes-Cantor genetic distance model and Neighbor-Joining method with 1000 bootstrap replications for consensus tree building.

### *FvJMT2* plasmid construction

The full-length *FvJMT2* C275 allele used to generate the overexpression construct (*35S:FvJMT2^C2^*^75^) was amplified from cDNA derived from *F. vesca* accession ES10 ^3^^9^ . The *35S:FvJMT2^A275^* construct was generated from *35S:FvJMT2^C275^*sequence using Q5® Site-Directed Mutagenesis Kit (New England Biolabs, USA). Finally, the truncated version of the A275 allele (*35S:FvJMT2^A275^*^Δ^*^C^*) was produced using the latter construct as a template. All CDSs were first cloned into pDONR entry vectors and then recombined into the pK7WG2D binary vector using the Gateway® Cloning system (Invitrogen, USA) under the control of the CaMV 35S promoter. The vector included also the fluorescent marker gene GFP expressed under the CaMV 35S promoter. Constructs were introduced into *Agrobacterium tumefaciens* strain GV3101 by electroporation. The GV3101 carrying the pK7WG2D empty vector was used as a negative control. All PCR amplifications were performed using iProof™ high- fidelity DNA polymerase (Bio-Rad Laboratories, USA), and all constructs were confirmed by Sanger sequencing. Primer sequences used for amplification and mutagenesis are listed in Table S3.

### Agroinfiltration of *FvJMT2* overexpression constructs in *Nicotiana benthamiana* and *Fragaria vesca*

Transient overexpression of the constructs carrying the different *FvJMT2* allelic variants, controls, and p19 was conducted in 3-week-old *N. benthamiana* leaves using a needleless syringe. Four days post-infiltration, GFP fluorescence was examined under a Leica DM2500 fluorescence microscope to confirm successful transformation. Transformed leaves were collected, immediately frozen in liquid nitrogen, ground to a fine powder, and stored at -80 °C until further processing. Similarly, *35S:FvJMT2^C275^* construct and the empty vector control were agroinfiltrated into immature green receptacles of *F. vesca* accessions ‘POR2’ and ‘POR3’. Agroinfiltrated fruits were monitored until ripening (∼10 days post-infiltration), harvested, immediately frozen in liquid nitrogen, and stored at −80 °C for subsequent analysis. Before grinding, the achenes were removed, and individual receptacles were ground to a fine powder.

*FvJMT2* overexpression was confirmed by quantitative PCR (qPCR). Total RNA was extracted from leaves and fruits using the Plant Total RNA Purification Kit (Norgen Biotek Corp., Canada), following the manufacturer’s instructions, and used for cDNA synthesis with the iScript™ cDNA Synthesis Kit (Bio-Rad Laboratories, USA). L23 60S was used as the housekeeping gene for *N. benthamiana*, and DBP for *F. vesca*. Leaves and fruits overexpressing *FvJMT2* were pooled to generate biological replicates for the subsequent biochemical analyses. Primer sequences are listed in Table S3.

### Methyl benzoate and methyl cinnamate quantification

Methyl benzoate and methyl cinnamate were quantified by GC-MS at the Institute for Plant Molecular and Cell Biology (IBMCP - Metabolomics Service, Valencia, Spain), as previously described ^85^. For *F. vesca* fruit ripening samples, volatile levels were quantified relative to those of the red (RR) stage. In the case of *Fragaria* species and agroinfiltrated *F. vesca* fruits, relative quantification was normalized to the average levels observed in *F. × ananassa* cv. ‘Camarosa’. In the case of *Fragaria* species and agroinfiltrated *F. vesca* fruits, relative quantification was normalized to the average levels observed in *F. × ananassa* cv. ‘Camarosa’. For *N. benthamiana* leaves, volatile levels in agroinfiltrated samples are expressed as peak area.

### Cloning, expression, and purification of recombinant FvJMT2 Protein

The full-length coding sequence of FvJMT2^C2^^75^ was cloned into pQE80L expression vector (Novagen, South Africa), which includes an N-terminal polyhistidine (6xHis) tag for affinity purification. The resulting plasmid was used to transform chemically competent XL1-Blue cells (Agilent Technologies, USA). Protein expression was induced by adding 0.2 mM isopropyl β-D-1- thiogalactopyranoside (IPTG), followed by incubation at 16 °C overnight. Recombinant proteins were purified using Ni-NTA Agarose (QIAGEN, Germany) according to the manufacturer’s instructions and concentrated using Amicon Ultra-30K centrifugal filter units (Sigma-Aldrich, USA). Protein concentration was determined using Pierce™ BCA Protein Assay Kit (Thermo Fisher Scientific, USA). Purified proteins were stored at −20 °C in 10 mM Tris-HCl, pH 8, 500 mM NaCl, 1 mM β-mercaptoethanol, and 30% glycerol. Protein integrity and equal loading were assessed by SDS-PAGE using 4–20% Mini-PROTEAN® TGX™ Precast Protein Gels (BioRad Laboratories, USA), followed by Coomassie Brilliant Blue (CBB) staining.

### Enzyme activity assay and kinetic analysis

Enzyme activity was assessed by measuring the transference of the [^14^C]-labeled methyl group from S-adenosyl-methionine (SAM) to the carboxyl group of various substrates: benzoic acid, cinnamic acid, salicylic acid, jasmonic acid, ferulic acid, (–)-quinic acid, *p*-coumaric, caffeic acid, gallic acid monohydrate, and *p*- hydroxyphenylacetic acid (*p*-HPAA). Radiochemical assays were performed in a total volume of 100 μL containing 50 mM Tris-HCl (pH 7.5), 2 mM substrate, 2.5 mM MgCl2, 5 mM dithiothreitol (DTT), 2.5 mM EDTA, 2 μL of [^14^C]-SAM (Perkin- Elmer, USA), and 30 μg of purified FvJMT2^C275^ protein. The reaction was initiated by the addition of SAM and incubated at 30 °C for 15 min (for benzoic acid, cinnamic acid, salicylic acid, and jasmonic acid), or 30 min for the rest of substrates. Reactions were stopped by the addition with 200 µL of 2-propanol supplemented with 1 M acetic acid. Then 300 µL hexane plus 150 µL Na2SO4 (6.7 %) were added, and the radioactivity of the upper organic phase was measured using a liquid scintillation counter (Beckman Coulter, USA).

For kinetic analysis, Michaelis-Menten constants (Km) were determined by varying the concentration of one substrate while the other was kept at a saturating level. The Km values were calculated by nonlinear regression fitting to the Michaelis-Menten equation and represent the average of three independent measurements.

### Drosophila suzukii colony maintenance

A laboratory colony of *D. suzukii* was maintained at the Instituto Valenciano de Investigaciones Agrarias (IVIA) under controlled environmental conditions (25 ± 1 °C, 60% relative humidity, 14:10 h light: dark photoperiod). Adults were reared in 25 × 25 × 25 cm BugDorm cages containing a 20% sugar solution and water-soaked cloth strips as continuous sources of moisture and energy. An artificial diet, based on cornmeal, sugar, brewer’s yeast, agar, and preservatives (methylparaben and propionic acid), was prepared weekly and distributed into 120 mL plastic vials, with 40 mL per vial, poured at an angle to increase the oviposition surface. Adult females oviposited directly on the diet, and the life cycle from egg to adult was completed in 13–15 days. Emerged adults were collected daily and transferred to new cages. Two BugDorms were maintained simultaneously and rotated monthly to ensure colony renewal and control of age structure.

### *D. suzukii* repellence/attractance assay

A two-choice behavioral assay was conducted using a 25 × 25 × 25 cm BugDorm cage (BugDorm-41515, MegaView Science Co., Ltd., Taiwan), featuring two plastic sides and four muslin mesh sides to allow for volatile diffusion. Inside the cage, four 50 mL plastic vials were randomly positioned: two contained control substrate (untreated strawberry purée), and the other two contained the same purée supplemented with the volatile compound under study at physiological concentrations. Strawberry purée was prepared from whole frozen strawberries (*Fragaria × ananassa*, Hacendado, Mercadona, Spain). 500 g of fruit were thawed at room temperature for 1 hour and then homogenized using a hand-held immersion blender (Braun Minipimer 5 MQ500 Soup, Germany) until a smooth and uniform consistency was obtained. For the assays, 30–50 mL of purée was dispensed into 50–150 mL plastic vials, providing sufficient headspace for flies to land on the inner walls and approach the purée surface. The vials were left uncovered to allow free access and olfactory detection. Volatile compounds were added to the purée at concentrations mimicking their natural levels in ripe *F. vesca* fruits, as reported by Urrutia et al. (2023)^38^. To this end, methyl benzoate (purity 99%, Sigma-Aldrich, CAS 93-58-3), methyl cinnamate (98%, Sigma-Aldrich, CAS 103-26-4) were each diluted in ethanol to prepare stock solutions. The appropriate volume of each stock was added directly at the time of mixing to 500 g of strawberry purée to achieve final concentrations of 196 ng/g fresh weight (FW) for methyl benzoate and 931 ng/g FW for methyl cinnamate. Control purées were prepared by adding the same volume of ethanol without any VOC. All purées were freshly prepared and used immediately for the behavioral assays. A cohort of 50 adult female *D. suzukii* (3–5 days old) was starved for 24 h prior to the experiment to enhance foraging motivation. Flies were released into the center of the cage, and the number of individuals found on each vial was recorded after 24 hours. Flies located elsewhere in the cage (i.e., not on the vials), were excluded from statistical analysis. All assays were conducted under controlled environmental conditions (25 ± 1°C, 70% relative humidity, 14:10 h light: dark photoperiod). Each experiment was replicated four times for each volatile compound tested.

## ACKNOWLEDGMENTS

This work was mainly supported by the European Research Council (grant number ERC Starting Grant ERC-2014-StG 638134), the Junta de Andalucía (grant numbers P20_00385 and UMA20-FEDERJA-115), and the Spanish Ministry of Science and Innovation and Universities (MICIU, PID2021-123677OB-I00) to D.P, and the Junta de Andalucía (UMA20-FEDERJA-093 and Postdoctoral program, and POSTDOC_21_00893) to C.M.-P. Further funding support was provided by the Margarita Salas Program from the Ministry of Universities - NextGenerationEU and Universidad de Granada (MS2021_70 to R.J.-M.), and by the Junta de Andalucía (grant numbers POSTDOC_20_00278 to M.U.). We thank Dr. José F. Sánchez Sevilla for facilitating growing the transgenic plants at the Andalusian Institute for Research and Training in Agriculture, Fishery, Food and Ecological Production (IFAPA) facilities in Churriana, Málaga, Spain, and Francisco Durán for its maintenance, respectively.

## CONTRIBUTIONS

R.J.-M. and M.U. conducted the characterization and functional validation of *FvJMT2*, and the GWAS analyses and identification of candidate genes, respectively. J.L.R. and A.G. performed the VOC quantification. T.T. and T.H. contributed to the *F. vesca* SNP panel. M.P.-H. and A.U. performed the behavioral assays with *D. suzukii.* J.J.S. contributed to the enzymatic characterization of FvJMT2. J.F.S. provided greenhouse facilities in Málaga (IFAPA). C.M.-P. and D.P. conceived, designed the research, and secured funding. R.J.-M., M.U., and D.P. wrote the article. All authors are involved in final manuscript editing

## SUPPLEMENTARY DATA

Supplementary Tables S1, S2, and S3 are available online.

## SUPPLEMENTARY FIGURES

**Figure S1.**
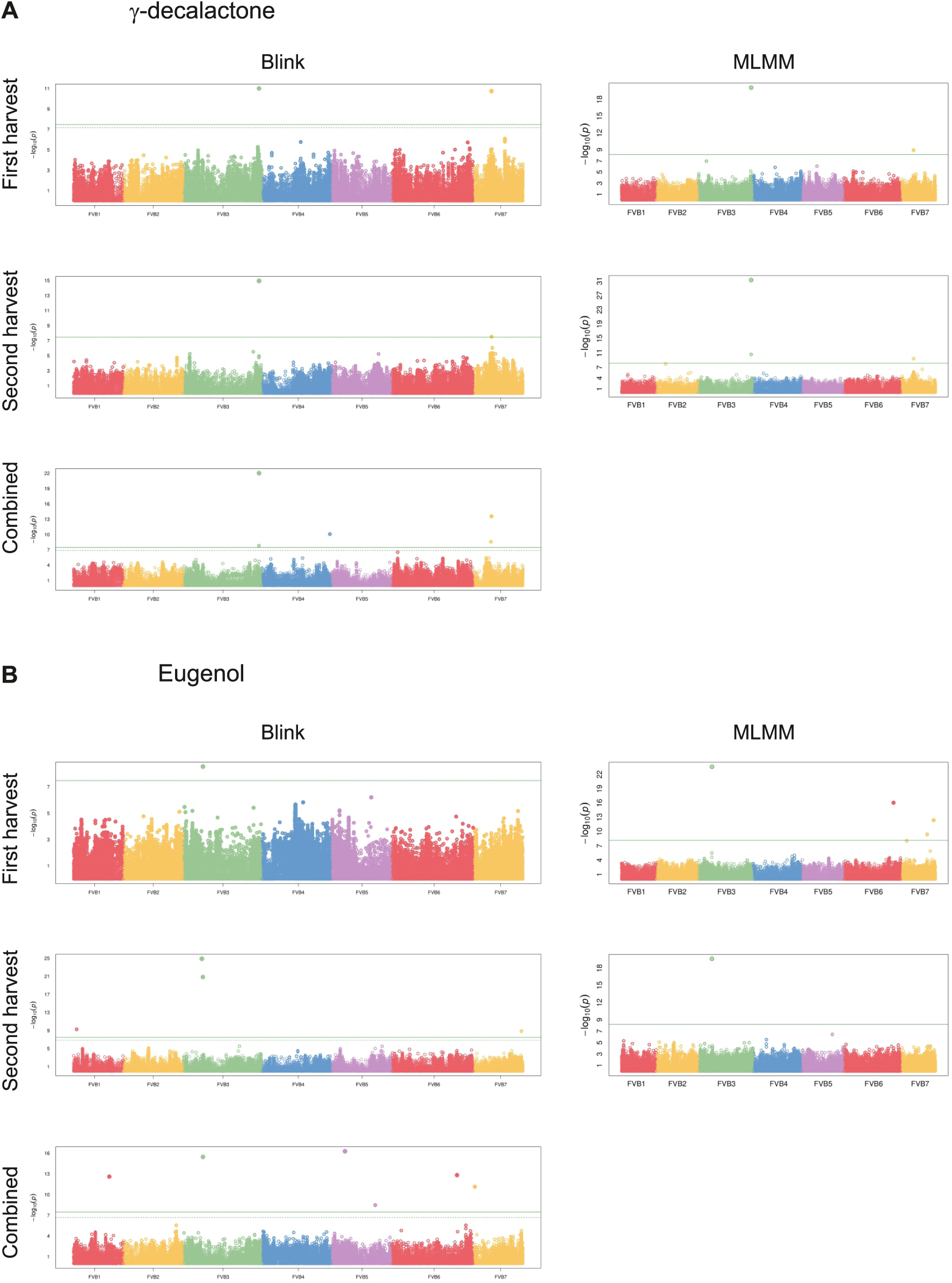
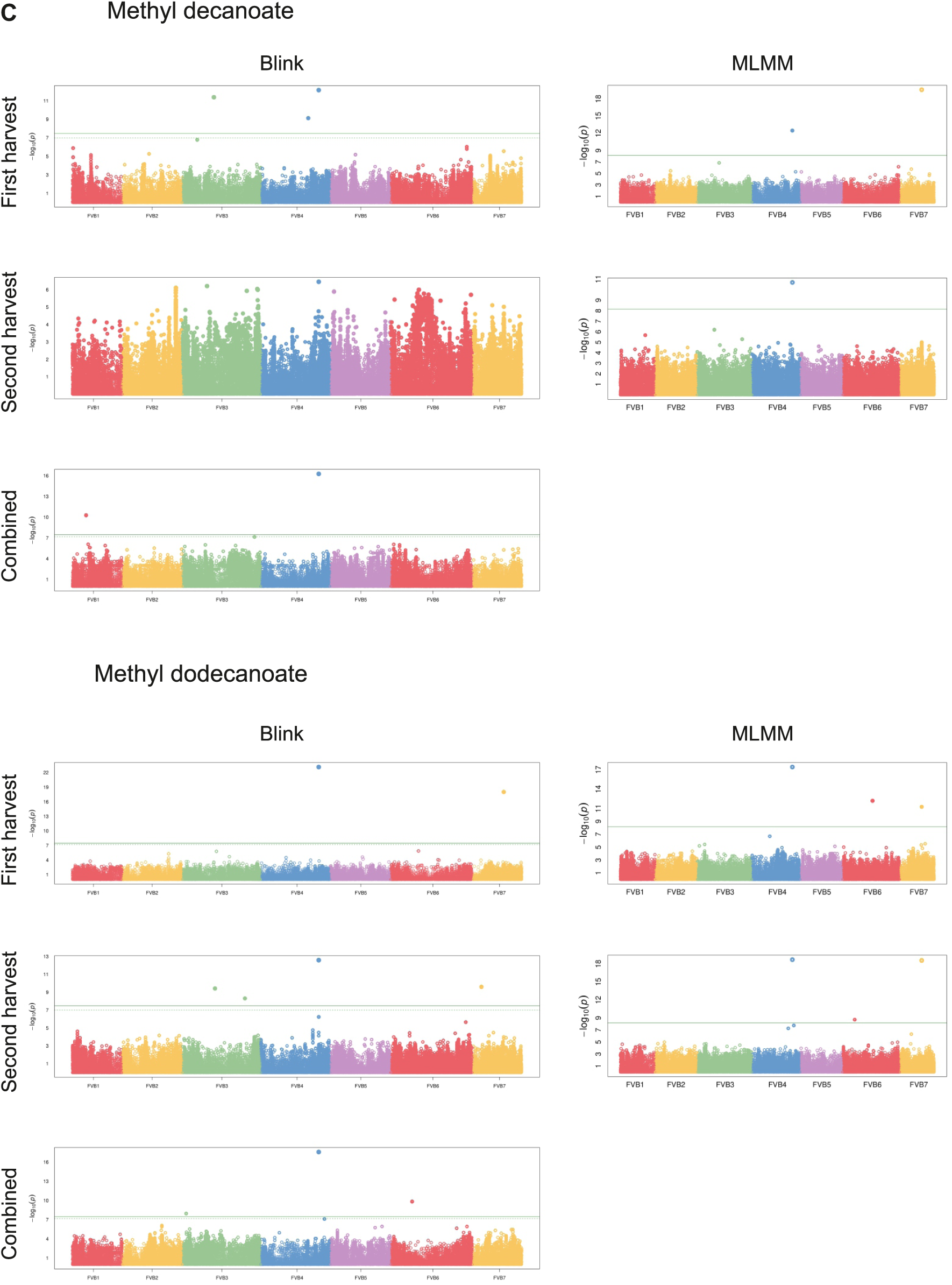
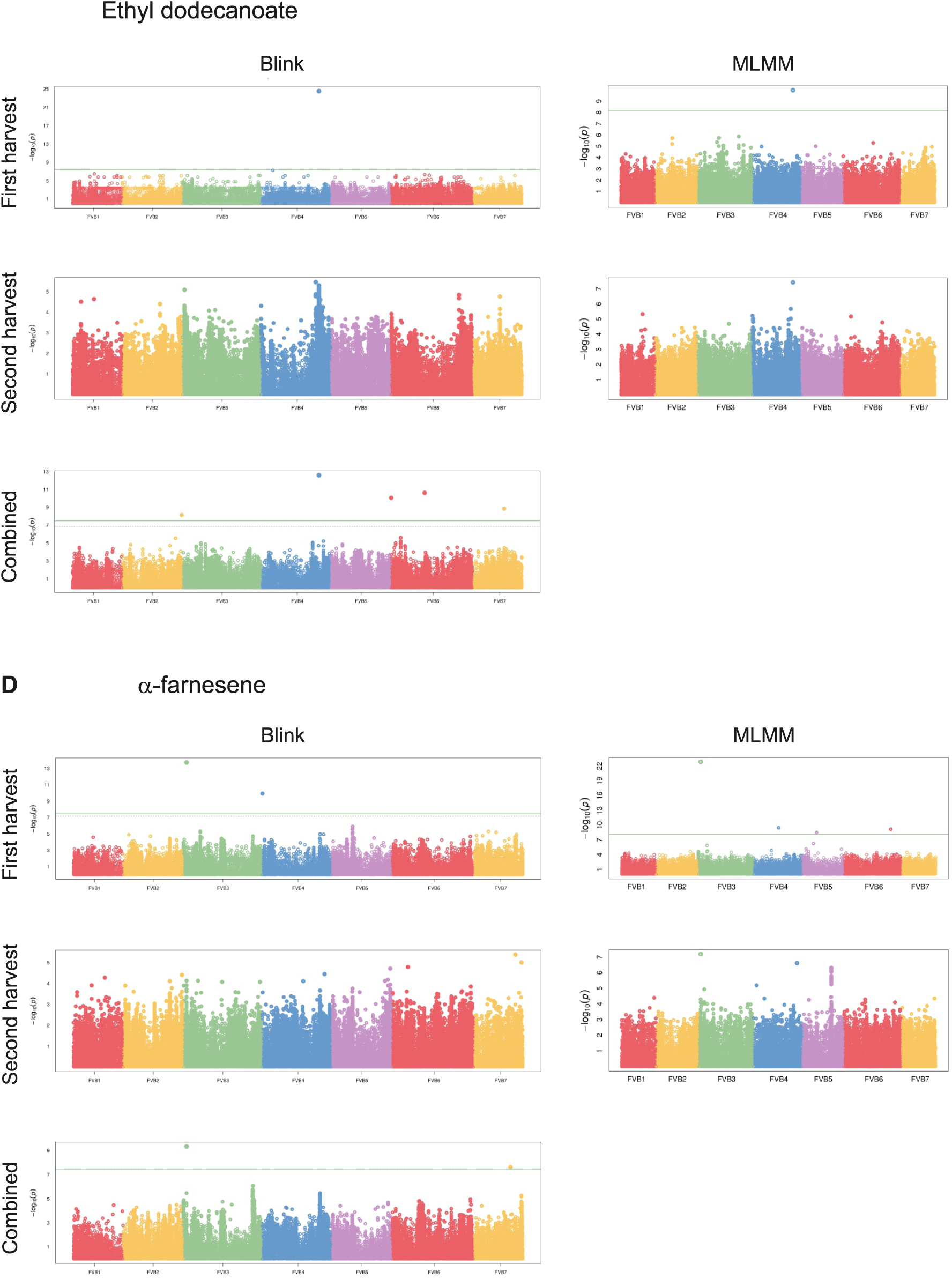
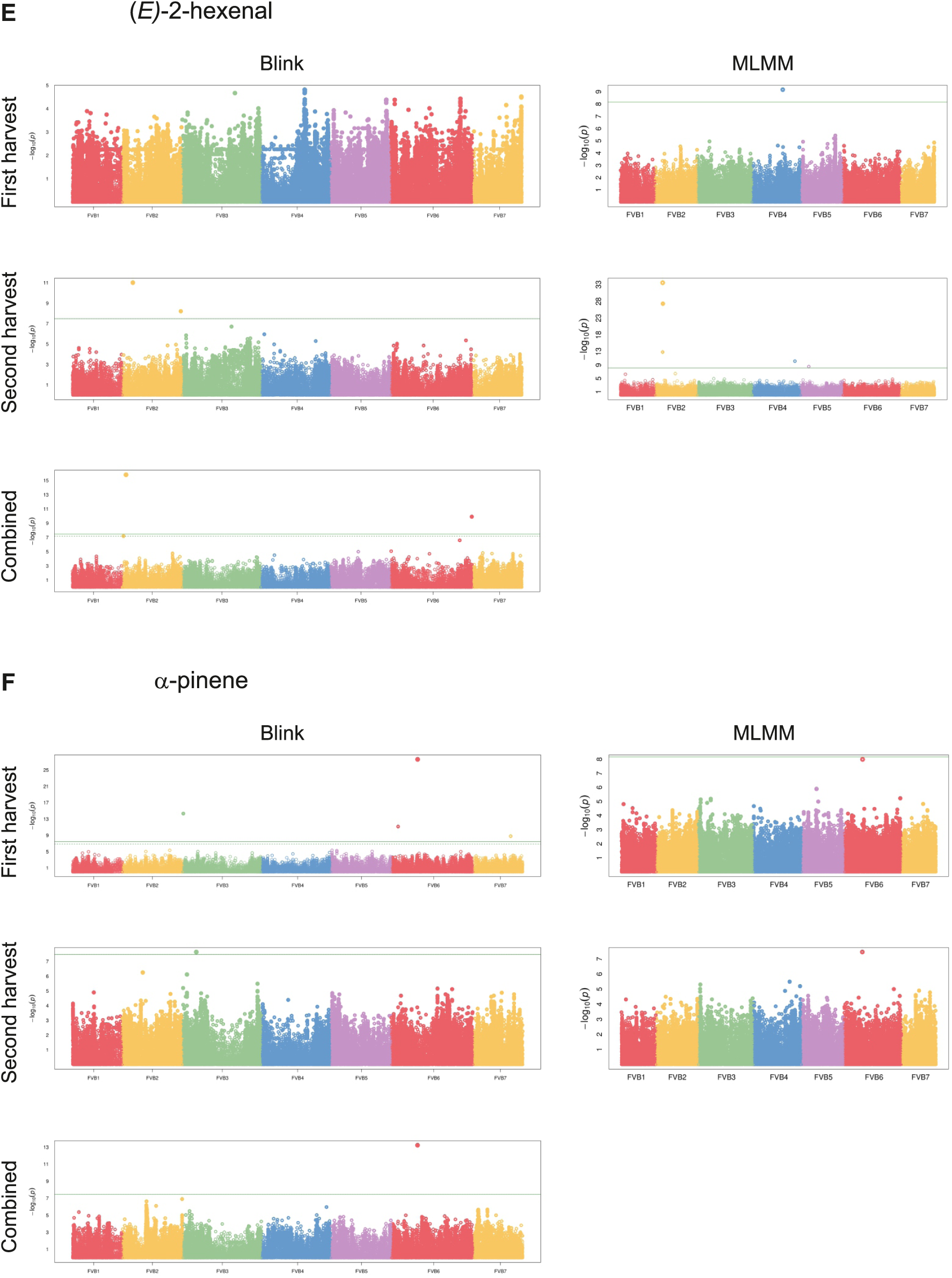
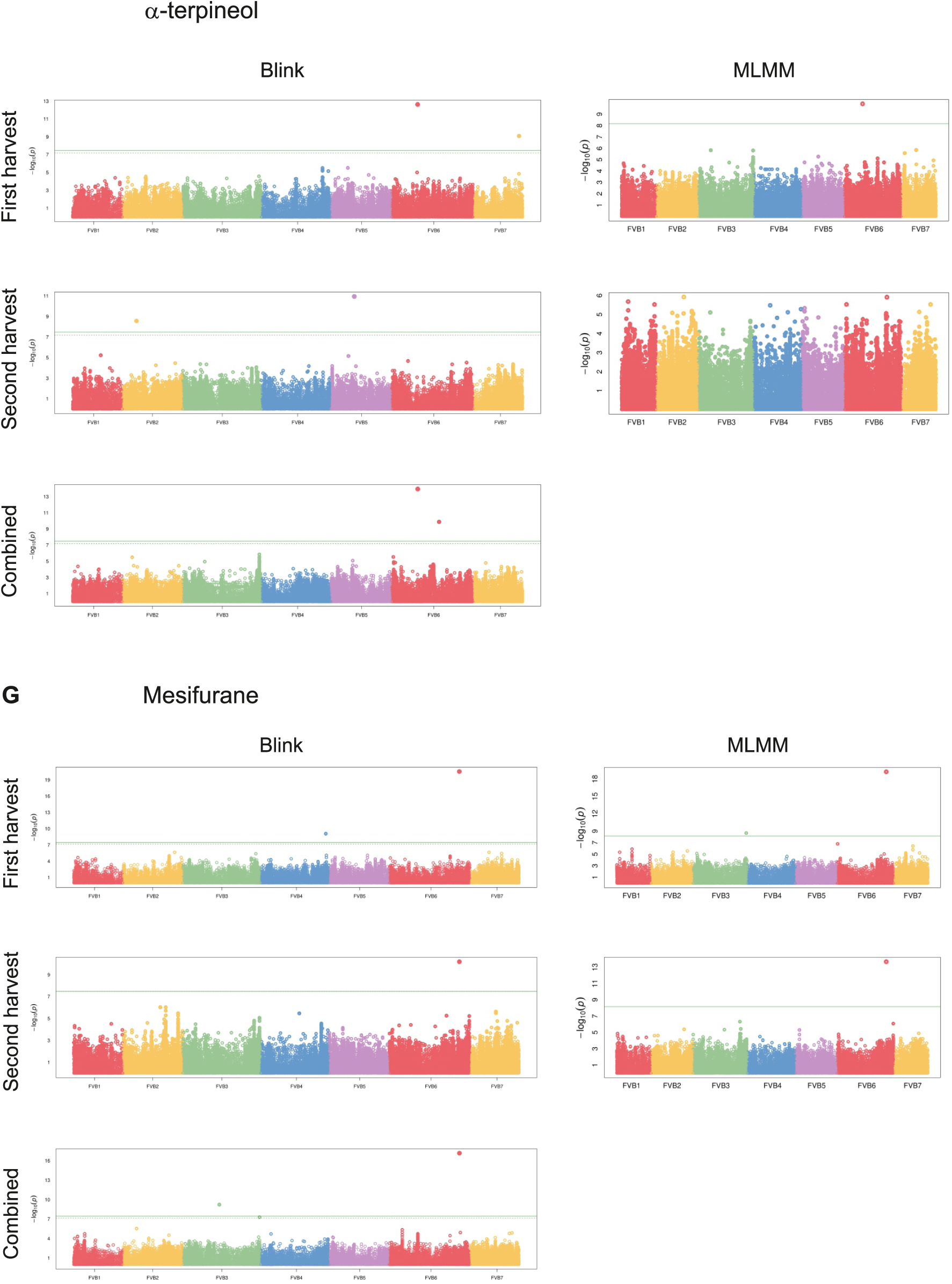
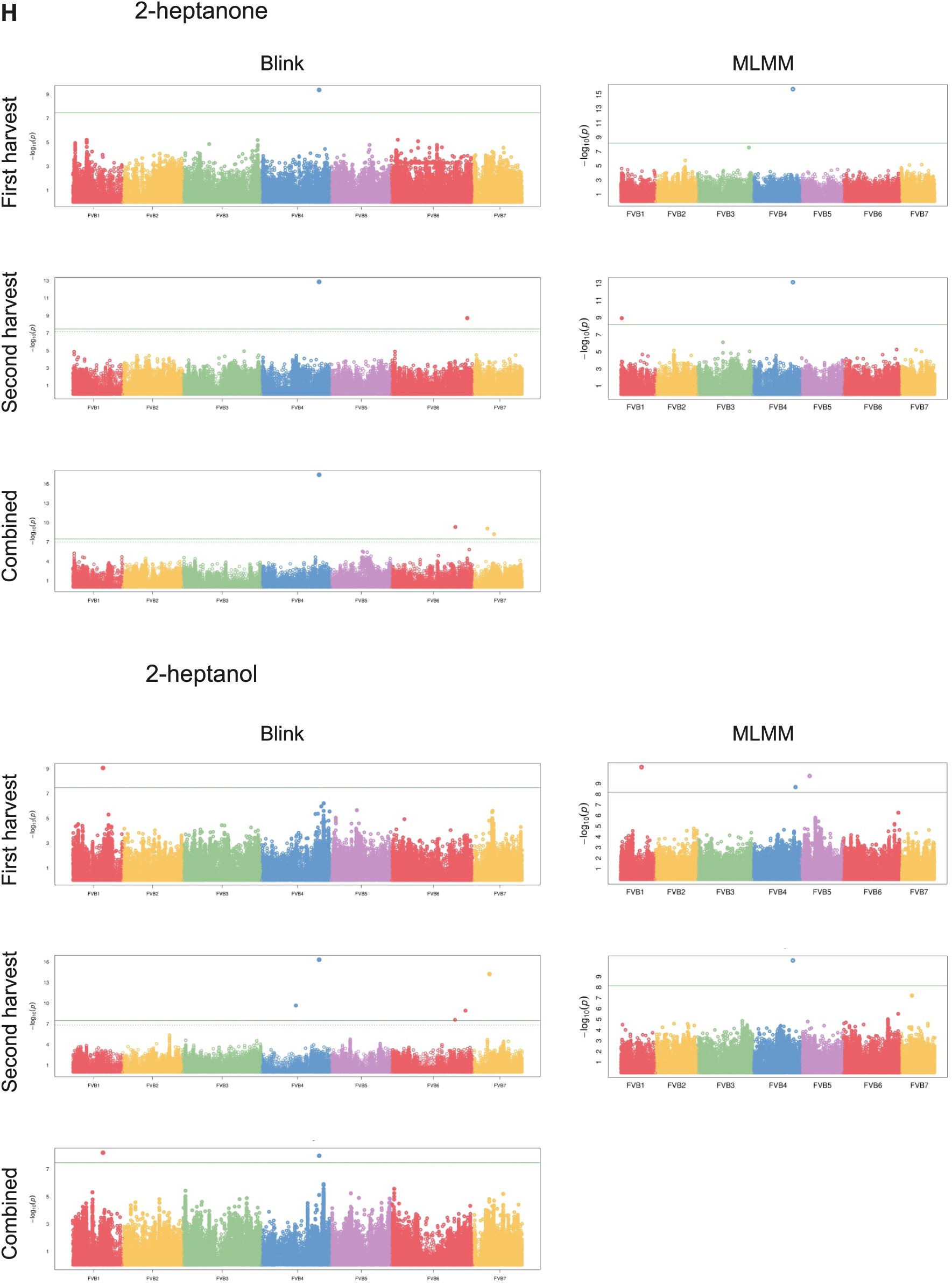
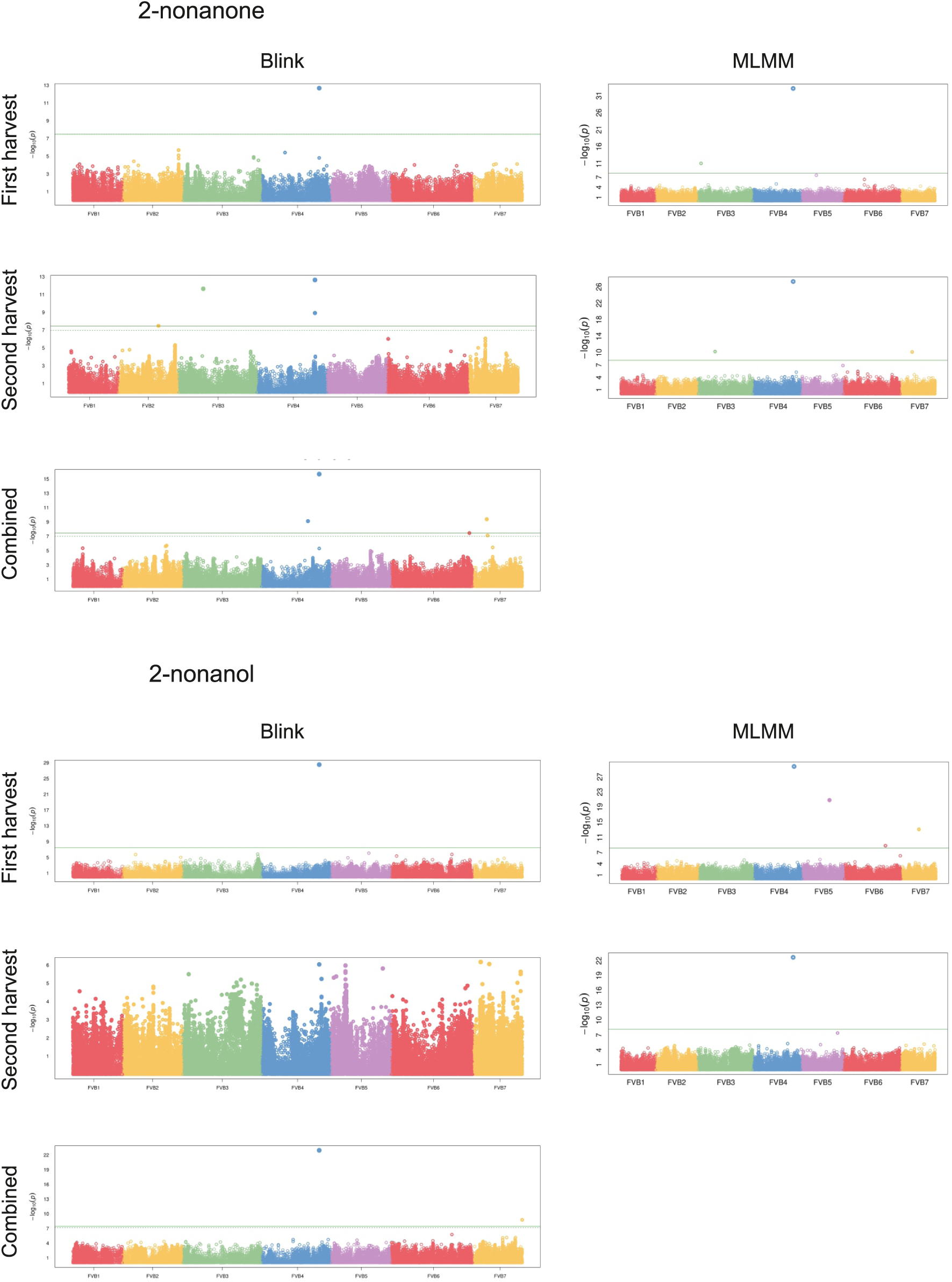
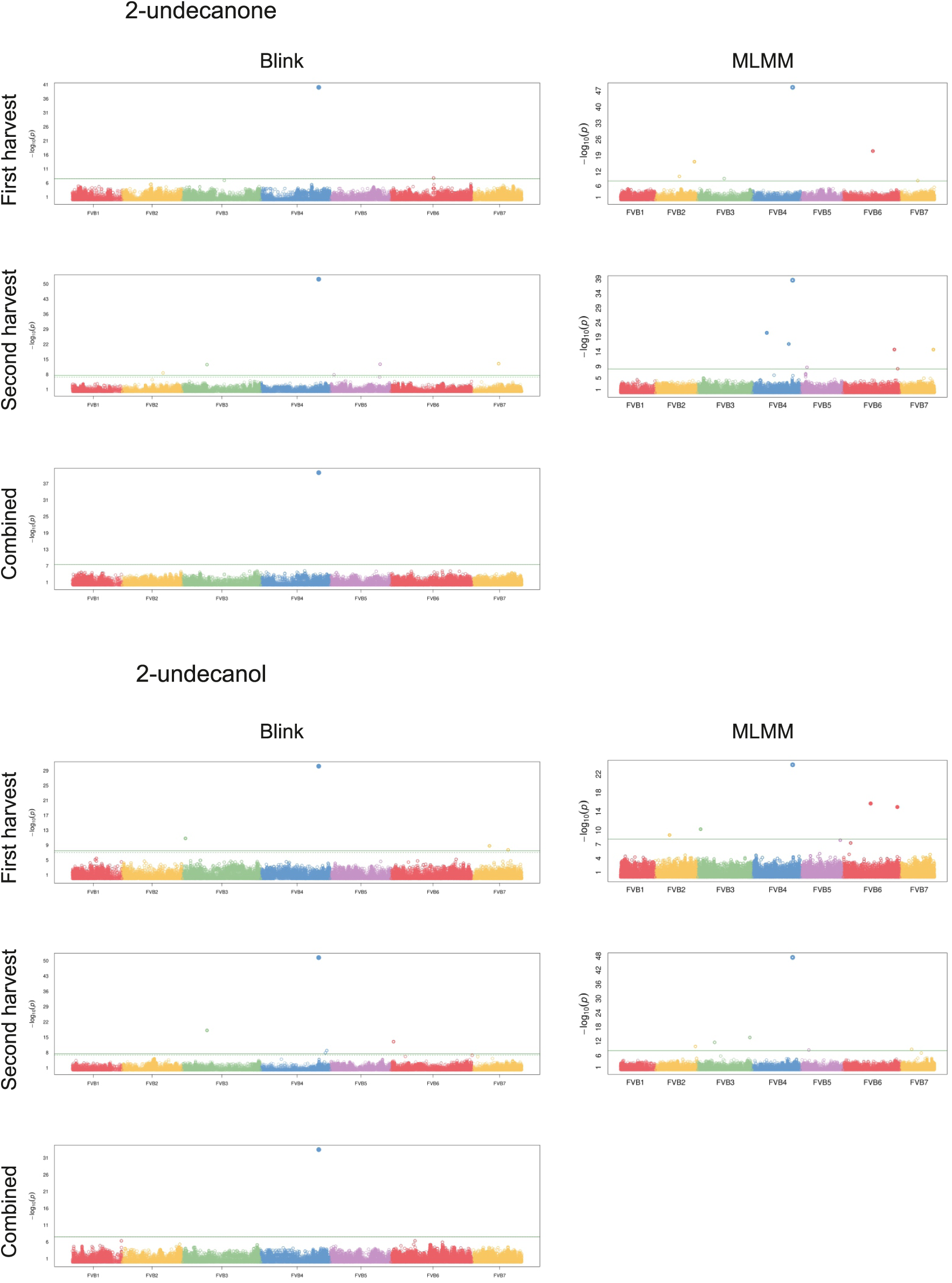
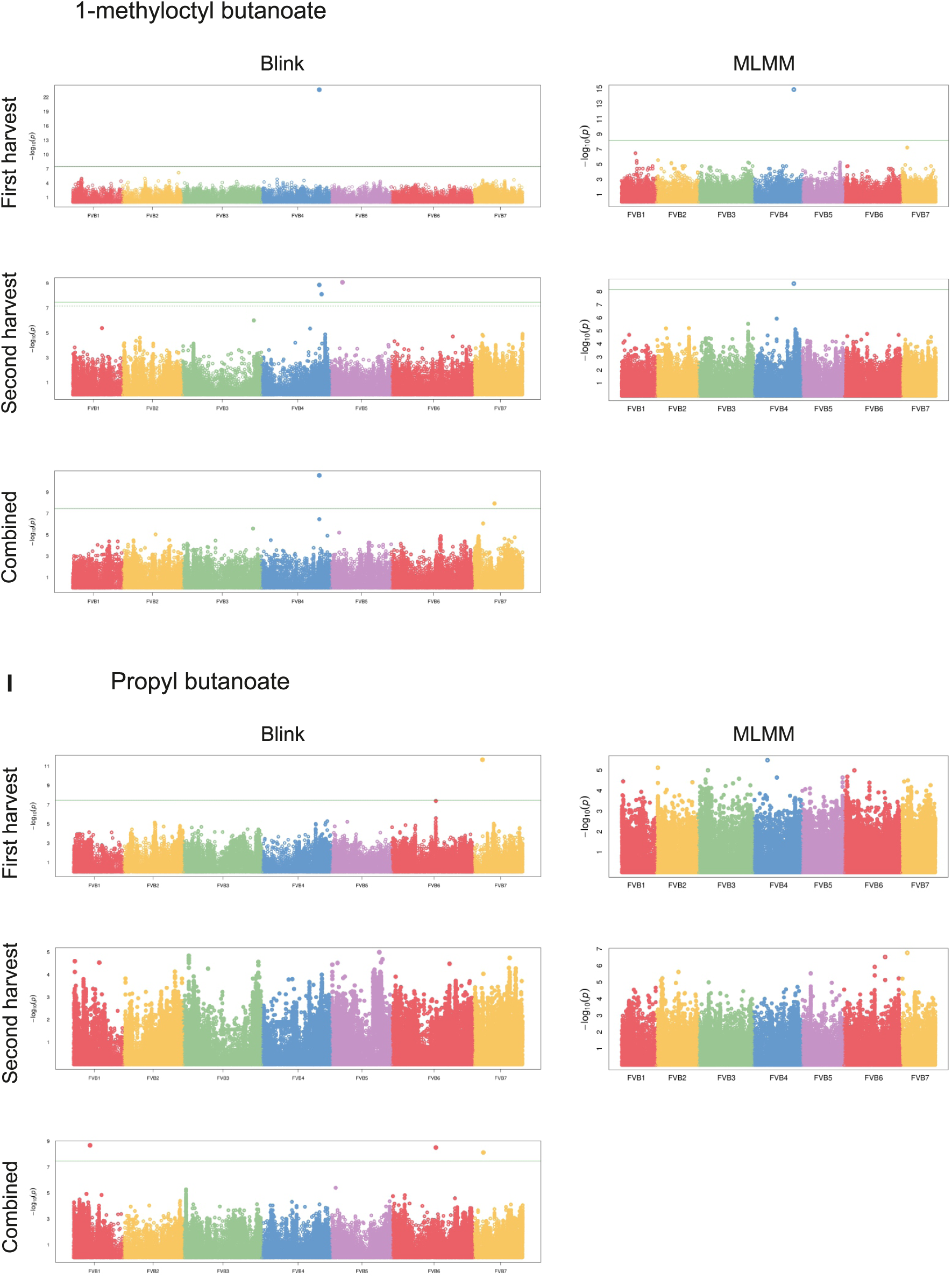
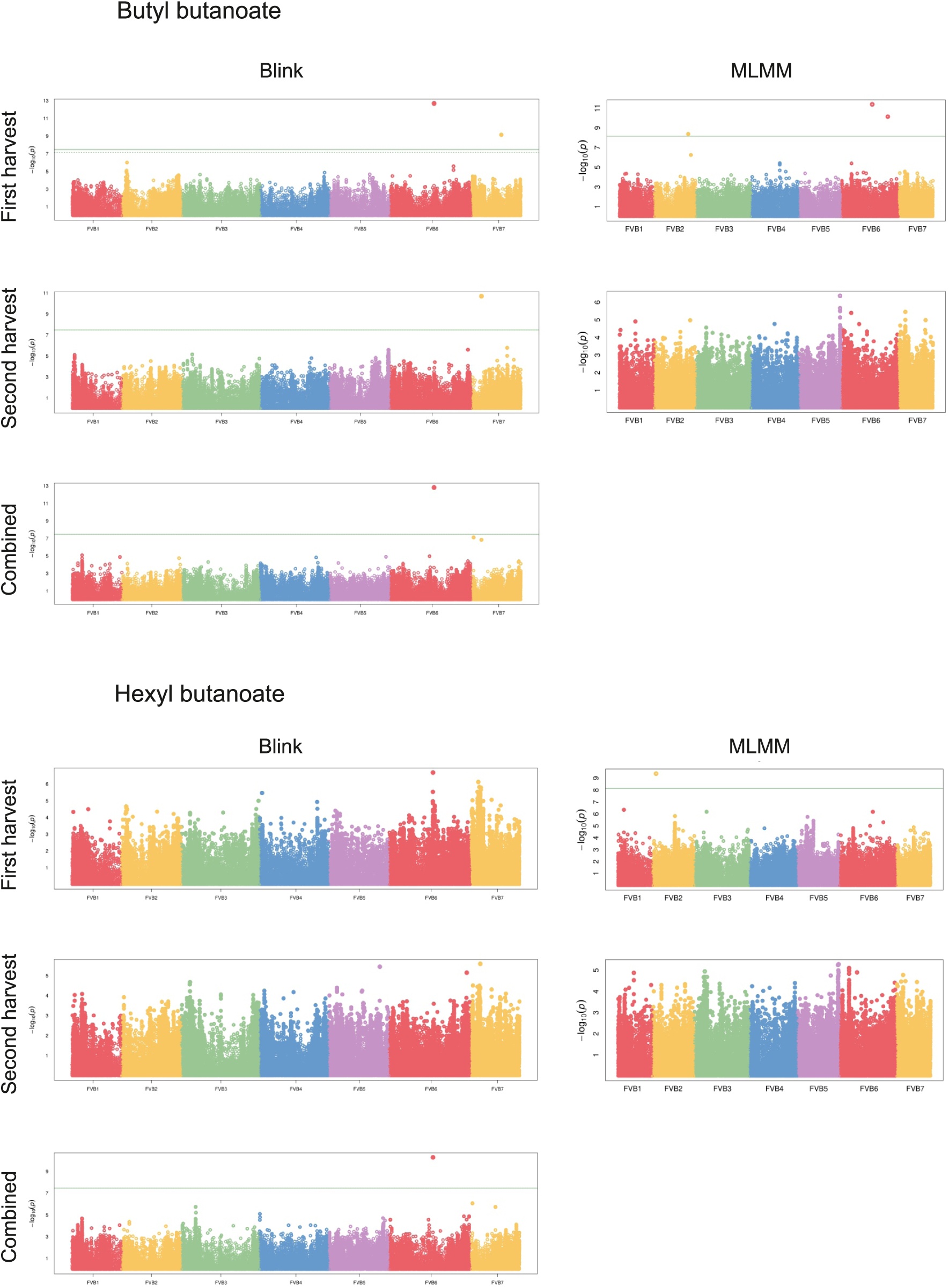
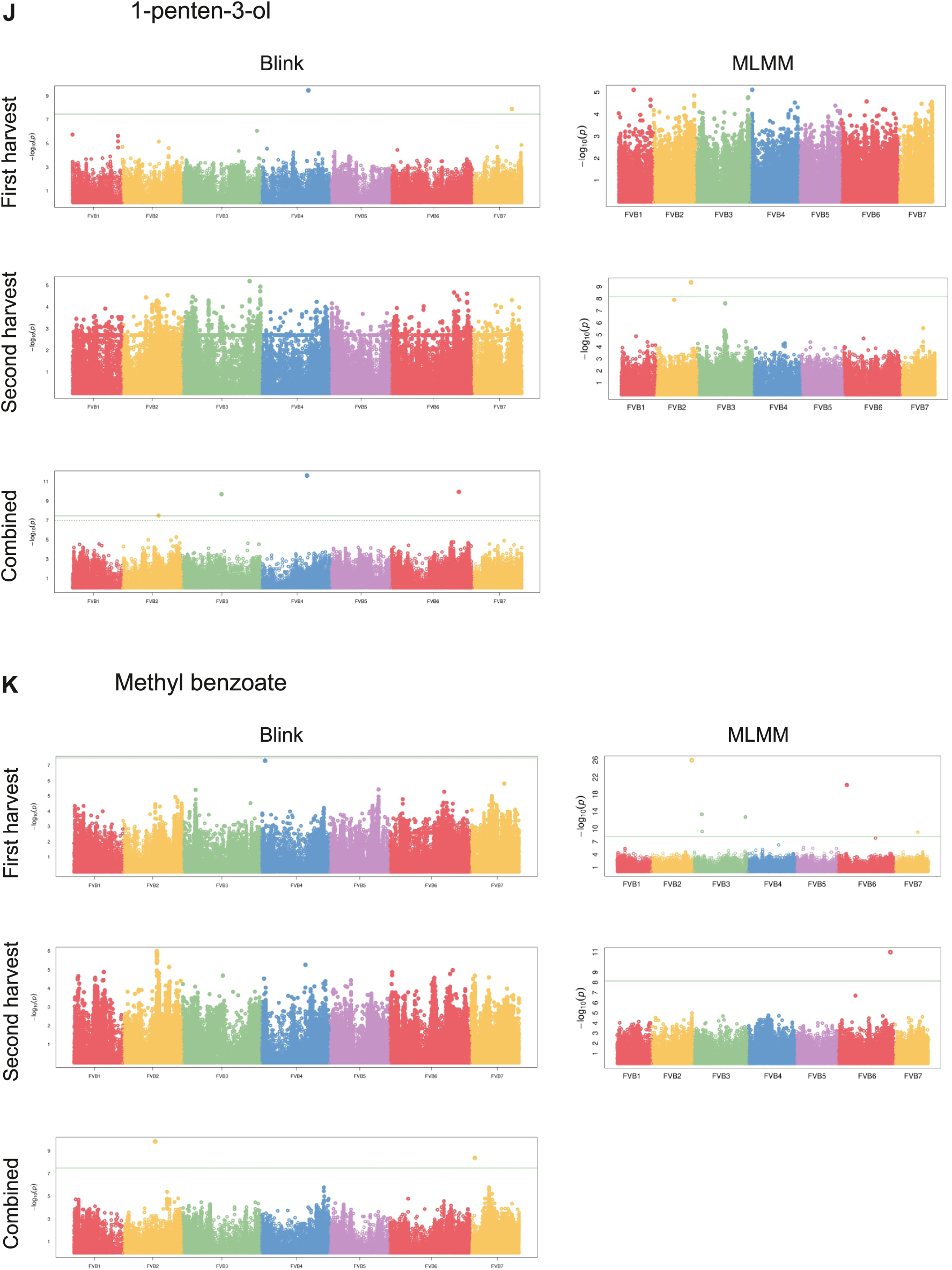
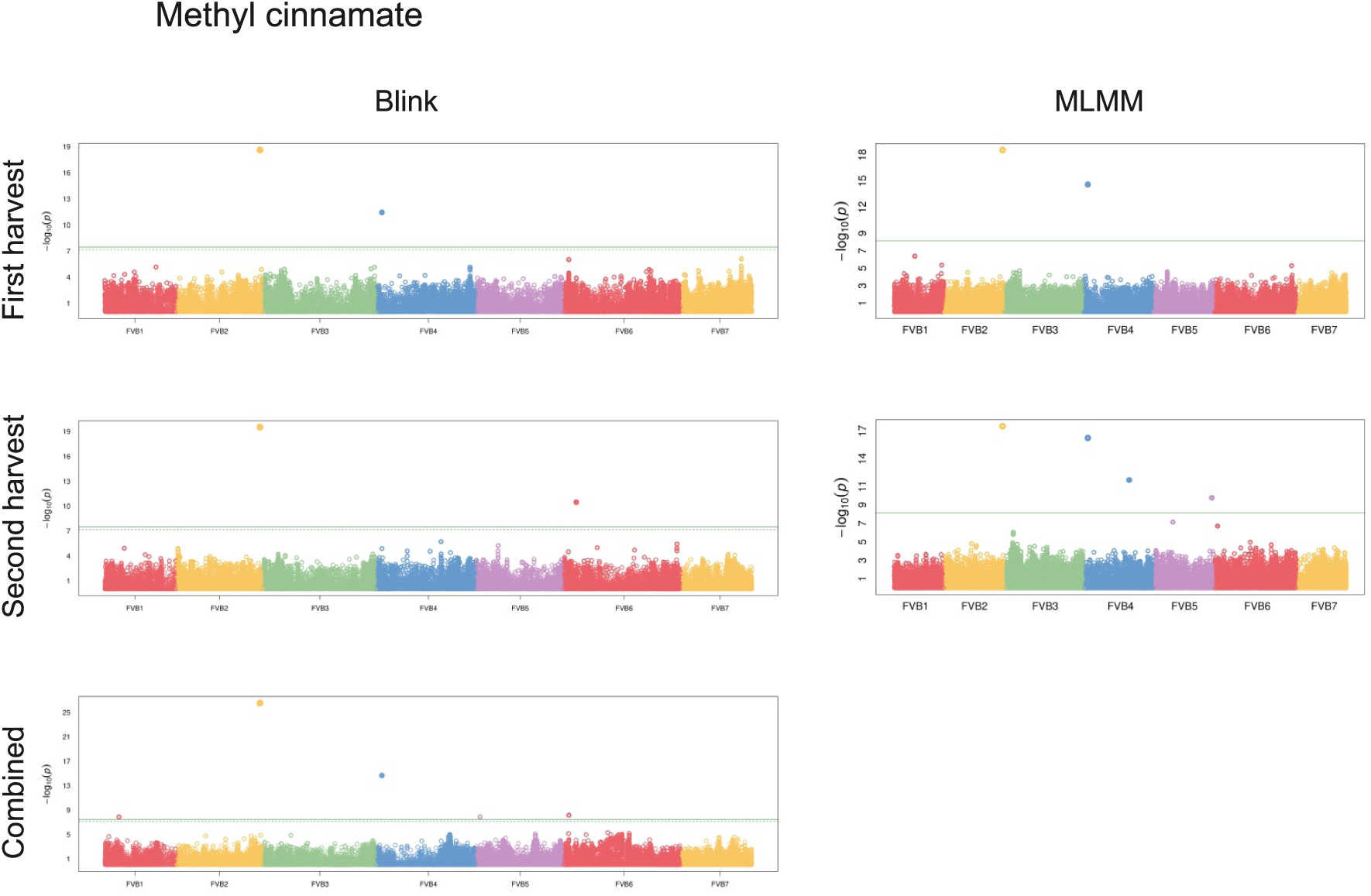
Manhattan plots of volatile compounds with stable GWAS associations. GWAS were conducted using the BLINK model for the first harvest, second harvest, and combined dataset (estimated marginal means), and the MLMM model for the first and second harvests only. The plots display volatile compounds for which consistent associations were detected in at least two datasets (individual harvests and/or combined dataset). Significant associations are shown for: γ- decalactone (A), eugenol (B), methyl decanoate, methyl dodecanoate and ethyl dodecanoate (C), (*E*,*E*)-α-farnesene (D), (*E*)-2-hexenal (E), α-pinene and α-terpineol (F), mesifurane (G), 2- heptanone, 2-heptanol, 2-nonanone, 2-nonaol, 2-undecanone, 2-undecanol and 1-methyloctyl butanoate (H), propyl butanoate, butyl butanoate and hexyl butanoate (I), 1-penten-3-ol (J), and methyl benzoate and methyl cinnamate (K).

**Figure S2.**
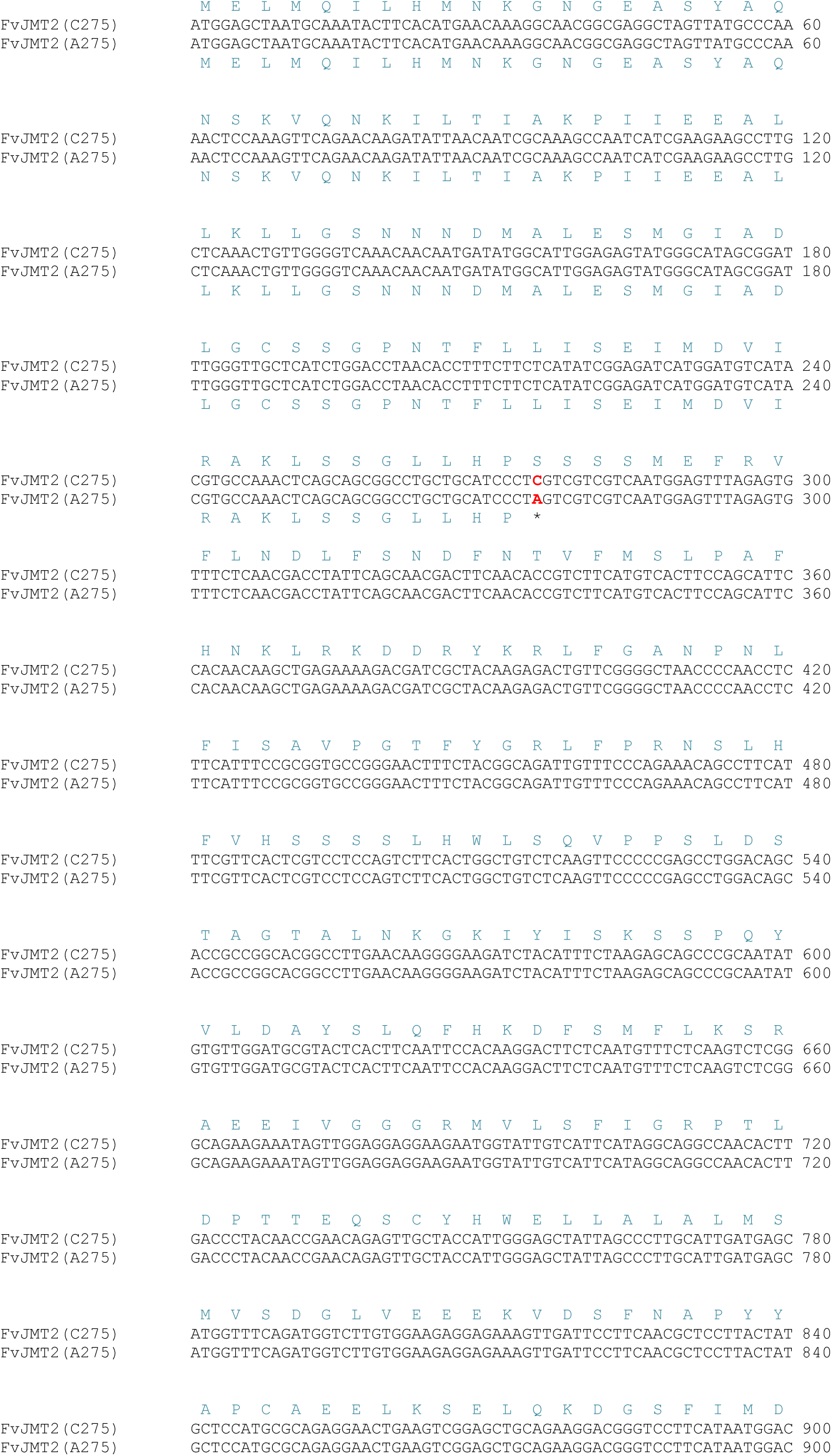
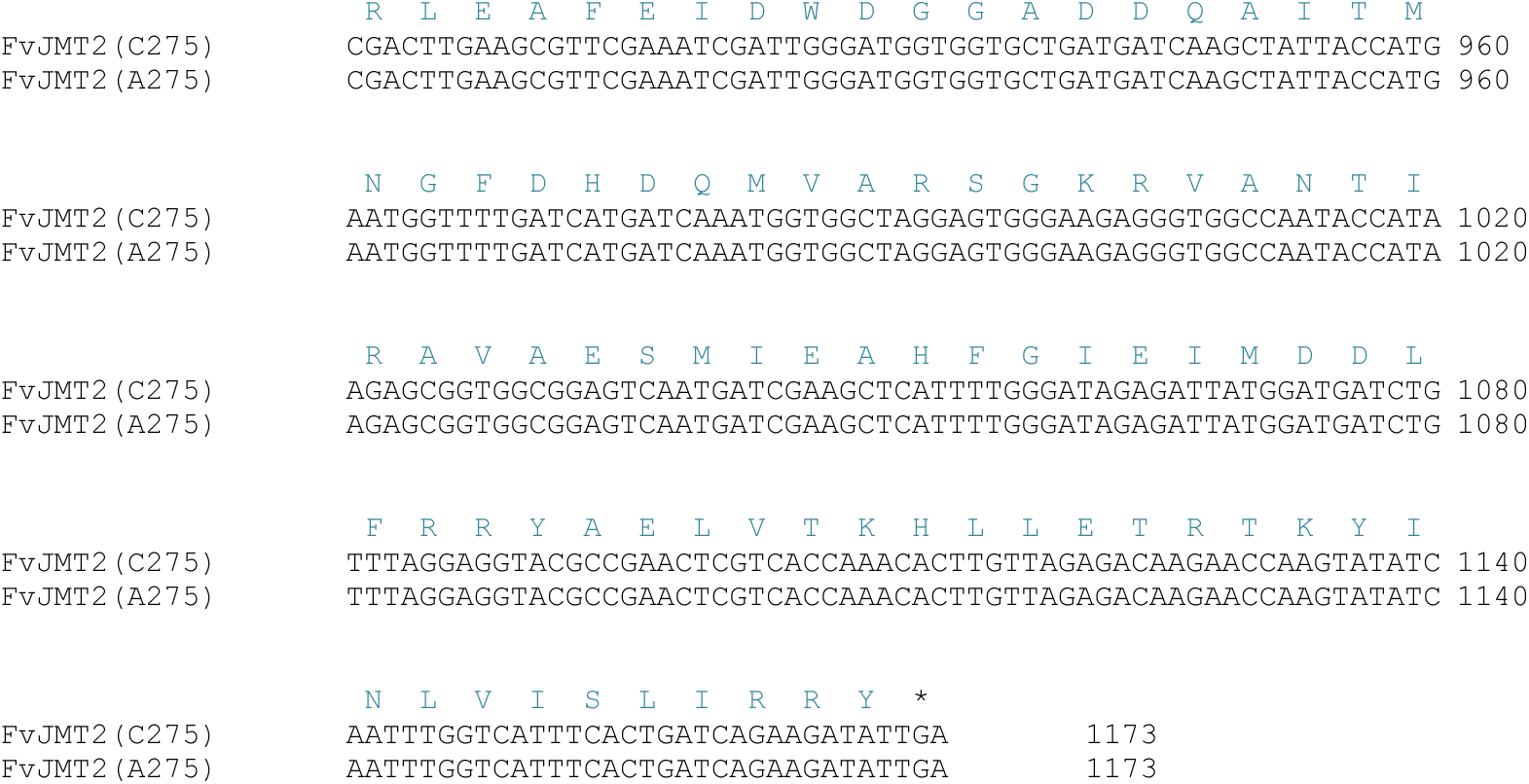
Alignment of FvJMT2 CDSs and protein sequences of the two alleles.

**Figure S3.**
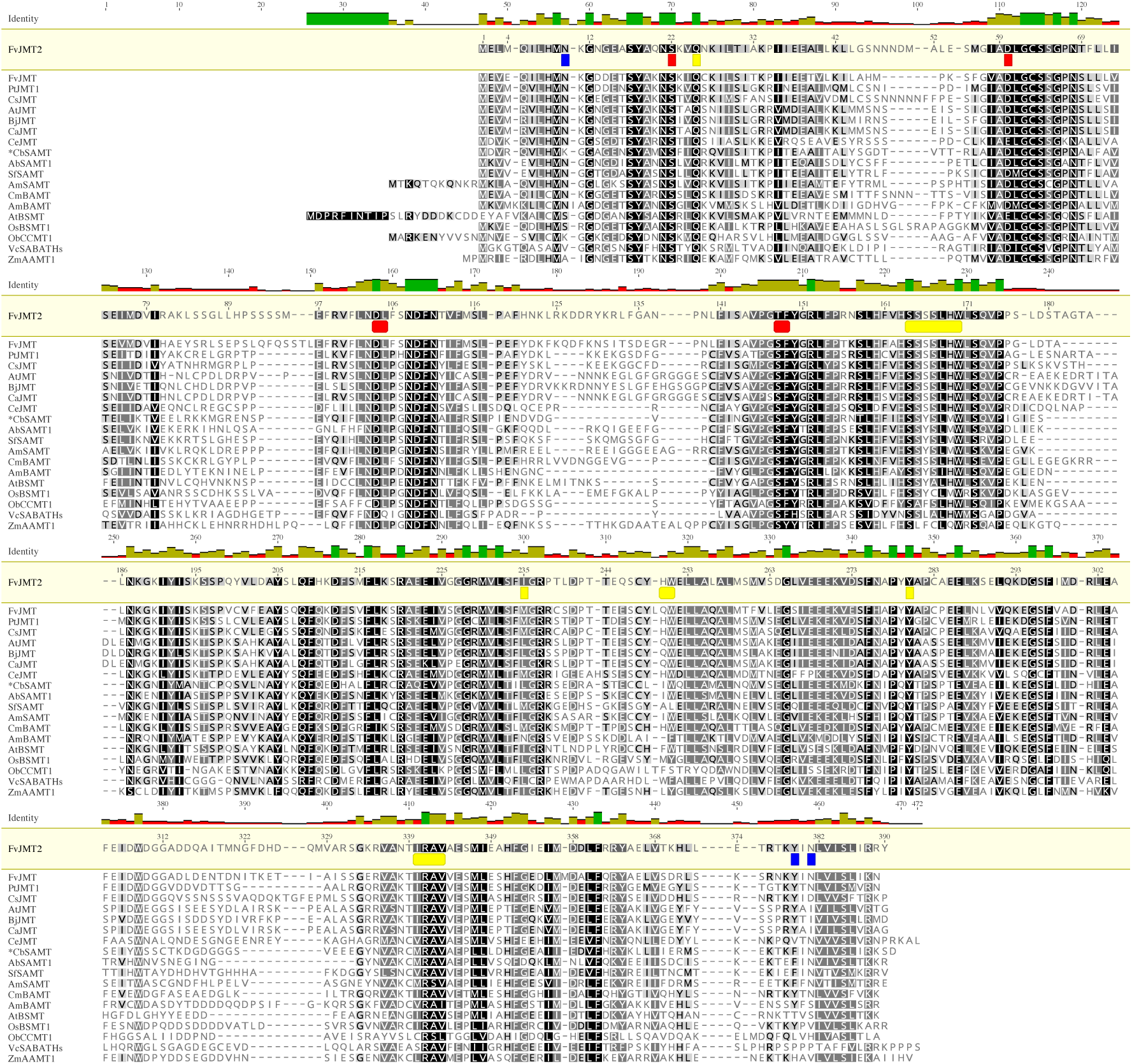
Multiple protein sequence alignment of FvJMT2 with other functionally characterized SABATH enzymes. Conserved amino acid residues corresponding to SAM/SAH (red), substrate- binding sites (yellow), and additional active sites (blue) are highlighted according to the enzymatic structure of CbSAMT (*) resolved by Zubieta et al. ^64^. Protein sequences were retrieved from GenBank (when available) or the corresponding publications: *Fragaria vesca* FvJMT ^35^, *Populus trichocarpa* PtJMT1 (KC894590), *Camellia sinensis* CsJMT (XP028087949), *Arabidopsis thaliana* AtJMT (AAG23343.1), *Brassica juncea* BjJMT (AY667499), *Capsicum annuum* CaJMT (DQ222856), *Cymbidium ensifolium* CeJMT (JQ360571), *Clarkia breweri* CbSAMT ^34^, *Atropa belladonna* AbSAMT1 (AB049752), *Stephanotis floribunda* SfSAMT (AJ308570), *Antirrhinum majus* AmSAMT (AF515284), *Cucumis melo* CmBAMT (XM_008468313.3), *Antirrhinum majus* AmBAMT (Q9FYZ9), *Arabidopsis thaliana* AtBSMT (BT022049*), Oryza sativa* OsBSMT1 (XM467504), *Ocimum basilicum* ObCCMT1 (EU033968), *Victoria cruziana* VcSABATHs (MZ541994), and *Zea mays* ZmAAMT (ZmAAMT1). Amino acid residues are color-coded based on their similarity to the FvJMT2 reference sequence: black = 100%, dark grey = 80–100%, light grey = 60–80%, and white < 60% similarity.

**Figure S4.**
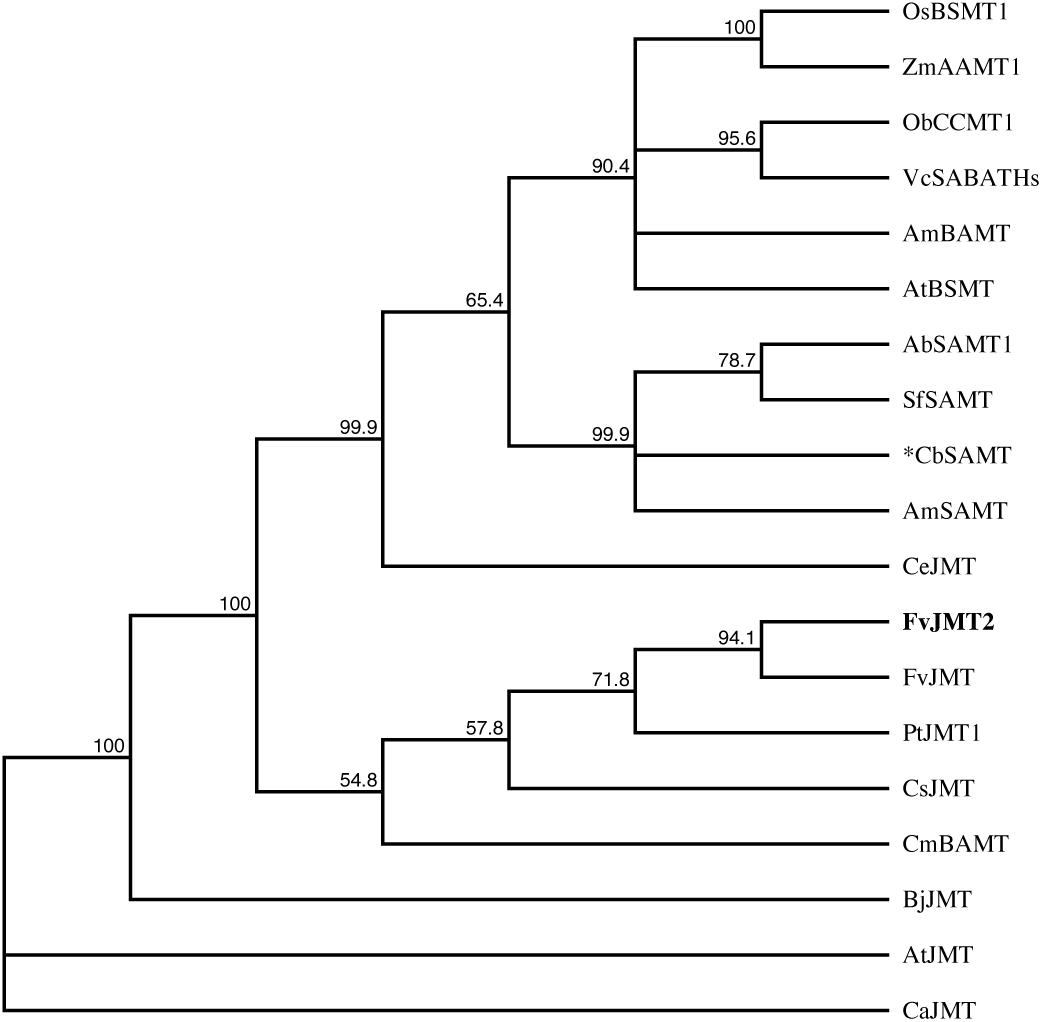
Neighbor joining consensus tree showing the relationship of FvJMT2 with other JMTs and CmBAMT. The phylogenetic tree groups FvJMT with jasmonic acid carboxyl methyltransferases (JMTs) from other plant species and with CmBAMT from *Cucumis melo* (*), supporting its functional and evolutionary similarity to these enzymes.

**Figure S5.**
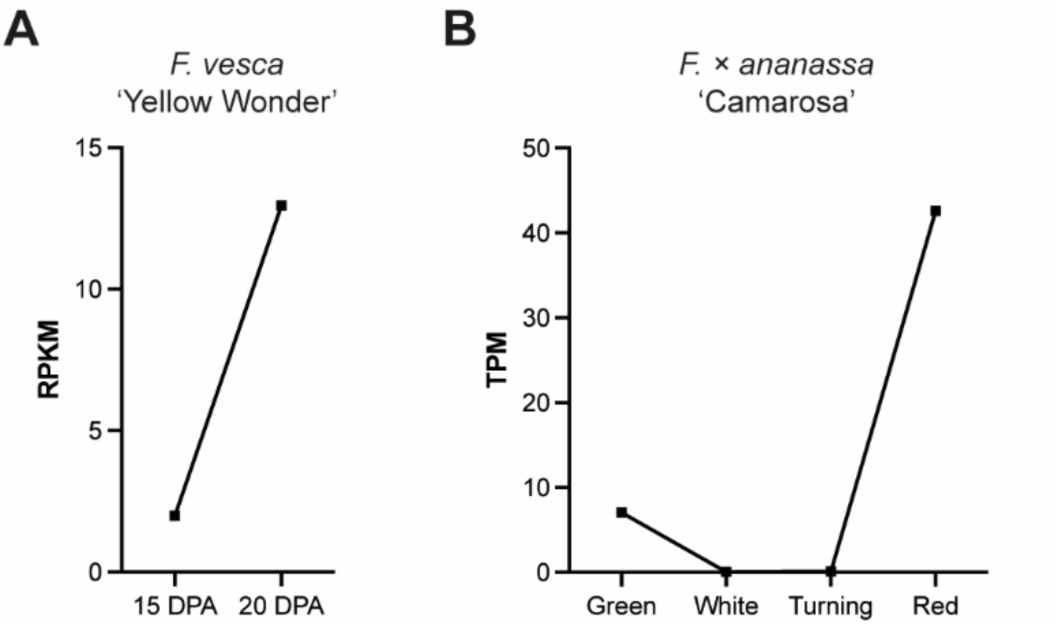
*JMT2* expression increases during strawberry fruit ripening in both diploid and octoploid genome backgrounds. **A)** Expression of *FvJMT2* at two timepoints (15 and 22 days post-anthesis, DPA) during fruit ripening in *F. vesca* ‘Yellow Wonder 5AF7‘, as reported by Hawkins et al. ^65^. **B**) Expression of the *F.* × *ananassa* ortholog *FaJMT2* (FxaC_8g06390) in the fruit receptacles at four developmental stages: immature green (Green), mature white (White), mature breaking (Turning), and mature ripe (Red), as previously reported ^61,62^.

**Figure S6.**
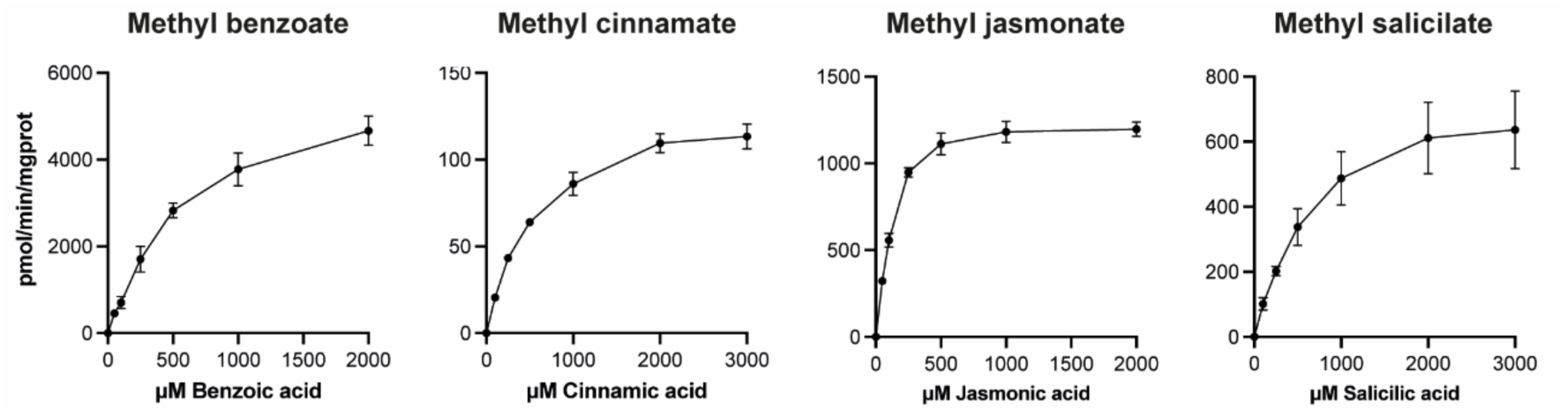
*In vitro* enzymatic kinetics of FvJMT2^C275^ with four major substrates. Michaelis-Menten curves showing the enzymatic activity of FvJMT2^C275^ in the presence of increasing concentrations of benzoic acid, cinnamic acid, jasmonic acid, and salicylic acid. Data represent the mean values of three independent assay replicates.

